# Reference glycan structure libraries of primary human cardiomyocytes and pluripotent stem cell-derived cardiomyocytes reveal cell-type and culture stage-specific glycan phenotypes

**DOI:** 10.1101/753707

**Authors:** Christopher Ashwood, Matthew Waas, Ranjuna Weerasekera, Rebekah L. Gundry

**Author notes:** CardiOmics Program, Center for Heart and Vascular Research; Division of Cardiovascular Medicine; and Department of Cellular and Integrative Physiology, University of Nebraska Medical Center, Omaha, NE, 68198, USA. Corresponding author: Rebekah L. Gundry, PhD University of Nebraska Medical Center Department of Cellular and Integrative Physiology 985850 Nebraska Medical Center Omaha, NE, 68198-5850, USA.

## Abstract

Cell surface glycoproteins play critical roles in maintaining cardiac structure and function in health and disease and the glycan-moiety attached to the protein is critical for proper protein folding, stability and signaling. However, despite mounting evidence that glycan structures are key modulators of heart function and must be considered when developing cardiac biomarkers, we currently do not have a comprehensive view of the glycans present in the normal human heart. In the current study, we used porous graphitized carbon liquid chromatography interfaced with mass spectrometry (PGC-LC-MS) to generate glycan structure libraries for primary human heart tissue homogenate, cardiomyocytes (CM) enriched from human heart tissue, and human induced pluripotent stem cell derived CM (hiPSC-CM). Altogether, we established the first reference structure libraries of the cardiac glycome containing 265 *N-* and *O-*glycans. Comparing the *N*-glycome of CM enriched from primary heart tissue to that of heart tissue homogenate, 21 structures significantly differed, and the high mannose class is increased in enriched CM. Moreover, by comparing primary CM to hiPSC-CM collected during 20-100 days of differentiation, dynamic changes in the glycan profile throughout *in vitro* differentiation were observed and differences between primary and hiPSC-CM were revealed. Namely, >30% of the *N*-glycome significantly changed across these time-points of differentiation and only 23% of the *N*-glycan structures were shared between hiPSC-CM and primary CM. These observations are an important complement to current genomic, transcriptomic, and proteomic profiling and reveal new considerations for the use and interpretation of hiPSC-CM models for studies of human development, disease, and drug testing. Finally, these data are expected to support future regenerative medicine efforts by informing targets for evaluating the immunogenic potential of hiPSC-CM and harnessing differences between immature, proliferative hiPSC-CM and adult primary CM.

## Introduction

In the heart, multiple cell surface glycoproteins, including calcium, potassium and sodium ion channels, work in concert to propagate electrical impulses to generate proper action potentials and subsequent contraction of the myocardium [2]. Cardiac glycoproteins are also required for maintaining intracellular substrate concentrations [3] and responding to perturbations in homeostasis [4]. The appropriate abundance, distribution, and post-translational modifications of these glycoproteins are required for proper function. Specifically, the glycan-moiety attached to the protein is critical for proper protein folding, stability and signaling [5]. For example, genetic defects that impact ion channel glycosylation sites result in impaired heart function [6]. Enzymatic removal of cell-surface *N-*glycan structures has a pronounced effect on mouse heart electrophysiology [7] and distinct glycan profiles have been reported for rat neonatal and adult ventricles [8]. Further supporting the role of glycosylation in heart maturation, treatment with sialidase to remove only a specific terminal monosaccharide (sialic acid) from adult ventricles results in sodium channel gating characteristics more similar to neonatal ventricles [7]. In humans, congenital disorders of glycosylation (CDG) which affect the glycan biosynthetic pathway and therefore the surface presentation of these sugars, have been associated with heart impairment in 16% (21 of 133) of disorders, with 10 of these CDGs affecting protein glycosylation specifically [9]. Finally, the glycosylation status of cardiac disease biomarkers such as natriuretic peptides (atrial (ANP), brain (BNP)) affects their function, stability in circulation, detectability by antibody-based methods, and interpretation of their clinical utility as it relates to specificity of disease [10–14].

Despite mounting evidence that glycan structures are key modulators of heart function and must be considered when developing cardiac biomarkers, we currently do not have a comprehensive view of the glycans present in the normal human heart. Ultimately, to understand the specific biological roles of glycoproteins, it is often necessary to define the population of each molecular species that arises from combinations of site-specific glycan diversity (*i.e.* microheterogeneity) at multiple sites of glycosylation, as this site-specific glycosylation may promote or interfere with interactions and signaling [1]. Moreover, in a complex organ like the heart, cell type specificity is a critical piece of the puzzle, as the context and directionality of glycan-mediated events defines, for example, how cells propagate action potentials [15] and multiple cell types interact during human development to properly form tissues [16]. Hence, defining the structures produced by the human cardiomyocyte protein glycosylation pathway, for example, could provide specific targets for studying glycogenes involved in regulating cardiomyocyte development, electrophysiology, disease biomarkers, and regenerative medicine efforts.

While rodent models have the practical advantage of providing accessible source material for research applications, they do not accurately represent human protein glycosylation owing to two specific enzymes: CMAH and GGTA1. The lack of CMAH in humans shifts the monosaccharide composition to Neu5Ac rather than Neu5Gc, although Neu5Gc can still be obtained through diet [17]. The lack of GGTA1 in humans means that this biosynthetic pathway of terminal di-galactosylation is absent in humans and cells featuring the glycan motifs produced by GGTA1 are immunogenic [18]. In both cases, non-human glycan structures found in animal models may confound studies of protein glycosylation of human cells. For these reasons, human source material, including primary cardiac tissue and stem cell derivatives, are most appropriate for generating reference glycan structure libraries to support future studies of glycoproteins within the context of human cardiac biology.

Modern mass spectrometry (MS) approaches enable the systematic discovery, characterization, and quantitation of glycopeptides and glycans. Specifically, methods for intact glycopeptide analysis by MS are becoming more readily available [19]. However, while native glycopeptides can be analyzed by these techniques, glycans must be released from their protein backbone for glycan structural isomers to be individually quantified. Moreover, the analysis of intact glycopeptide data requires that the search space be defined *a priori,* including both protein and glycan databases [20]. Accurate databases that reflect glycan compositions present within a sample are critical for sensitivity (probability of a correct glycopeptide match to a spectrum) and specificity (probability of an unassigned spectrum given that the spectrum is not actually a glycopeptide), as a glycan composition must be within the search space for corresponding glycopeptides to be identified. While glycans released from homogenized human heart tissue have been profiled using MALDI-MS [21], this approach does not resolve structural isomers or provide cell-type specific information.

In the current study, we used porous graphitized carbon (PGC) LC interfaced with MS (PGC-LC-MS), a technique that resolves glycan structures and enables characterization and quantification of individual structures [22], to generate glycan structure libraries. By applying this approach to primary human heart tissue homogenate, cardiomyocytes (CM) enriched from human heart tissue, and human induced pluripotent stem cell derived CM (hiPSC-CM), we established the first reference structure libraries of the human cardiac glycome containing 265 *N-* and *O-* glycans. These data will benefit future functional glycomics studies of CDG as the glycan structural data will inform lectin- and glycan-array mapping approaches by defining the end-result structures that are the result of responsible genes, proteins and metabolites. The glycan isomer libraries defined in this study are necessary for interpretation of future intact glycopeptide data and the cell-type specific glycan differences observed here will support the generation of cell-type specific protein glycoform maps of the human heart, thereby promoting the development of cell type-specific targeting strategies. Moreover, by comparing primary CM to hiPSC-CM collected during 20-100 days of differentiation, dynamic changes in the glycan profile throughout *in vitro* differentiation and differences between primary and hiPSC-CM are revealed. These observations are an important complement to current genomic, transcriptomic, and proteomic profiling and reveal new considerations for the use and interpretation of hiPSC-CM models for studies of human development, disease, and drug testing. Finally, these data are expected to support future regenerative medicine efforts by informing targets for evaluating the immunogenic potential of hiPSC-CM and harnessing differences between immature, proliferative hiPSC-CM and adult primary CM.

## Materials and Methods

### Study design

An overview of the study design is shown in Figure 1. A comparison of CM enriched from human cardiac tissue to matching tissue homogenate was used to identify glycan structures preferentially expressed in CM over other cell types in the heart. Cadaverous heart tissue from three donors (2 male, 1 female) were used for this study. For hiPSC-CM, cells were collected across 20 timepoints of differentiation to enable assessment of temporal differences. Finally, as hiPSC-CM culture includes eukaryotic cell culture components which may be glycosylated [23], we profiled the constitutive glycan structures in all cell culture reagents used and developed a method to confirm that these culture components do not directly contribute to the glycan profile of hiPSC-CM (Figure S4).

**Figure 1.**
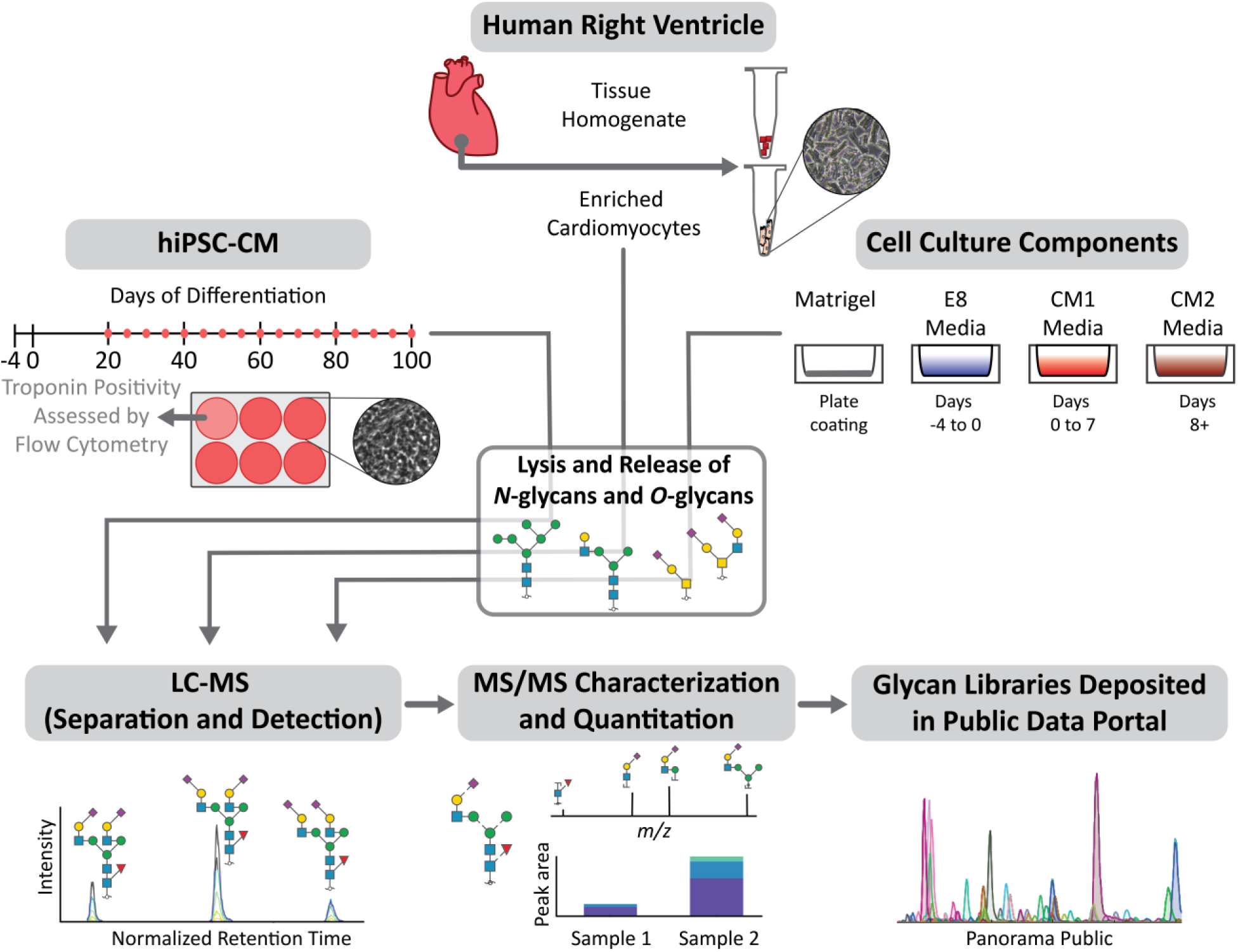
Schematic outline of the study design investigating protein glycosylation in human cardiomyocytes. Glycans from cardiomyocytes enriched from human heart tissue (n=3), matched tissue homogenate (n=3), hiPSC-CM collected across days 20-100 of differentiation (n=3 for each time-point), and hiPSC-CM culture components (n=1) were analyzed using PGC-LC-ESI-MS/MS for structure-based glycan characterization and quantitation. Glycan structure libraries are publicly available in Panorama to facilitate future glycoproteomic and glycomic efforts.

### Cardiomyocyte enrichment from human tissue and fixation for artifact identification

Frozen right ventricle myocardial tissues (500 mg) from three non-failing adult human donors (Table S1) were thawed at 37 °C for 5 min followed by dissection in 5 mL of Ca^2+^ free solution to ∼1 mm^3^ pieces. The tissue piece suspension was placed in a spinner flask (Integra Biosciences, Hudson NH) and washed 3x with warm Ca^2+^ free PBS solution. Tissue pieces were resuspended in 20 mL of collagenase solution, digested for 10 min at 37°C then supplemented by 10 mM CaCl_2_ followed by an additional 30 min of digestion at 37°C. The digested tissue pieces were then triturated and filtered (300 μm filter, pluriSelect, Leipzig, Germany). The filtrate was collected, resuspended in storage solution and centrifuged at 95 x *g* for 10 min. The pellet was collected and resuspended in PBS. A representative bright-field image of the cells collected by this approach is shown in Figure S1A, demonstrating that the vast majority of cells are CM. To verify that changes in glycan structure quantity are not artifacts of residual enzymatic activity during sample handling, a comparison of CM enriched by the above stated method were compared to CM enriched after tissue pieces were fixed with 4% paraformaldehyde for 2 hours, followed by the steps described above to enrich CMs (Figure S2).

### Cell culture and validation of cardiomyocyte identity in hiPSC-CM cultures

DF6-9-9T hiPSCs were maintained in monolayer culture and differentiation was performed as described [24], generating predominantly ventricular-like cells. Quality control (QC) evaluation of hiPSC-CM differentiation was performed by following the Standard Operating Procedure for flow cytometry-based assessment of troponin positivity within hiPSC-CM cultures as previously described [25] and experimental details are provided in the Supplementary Methods (Table S3). A representative flow cytometry histogram showing percent positivity for troponin (>85%) is shown in Figure S1B. hiPSC-CM were collected as a pellet (∼1x10^7^ cells) and frozen at -80°C until use.

### Sample preparation for glycan analysis

Primary enriched CM and hiPSC-CM were lysed in 1X Invitrosol (Thermo Fisher Scientific, IL, USA), 100 mM ammonium bicarbonate (NH_4_HCO_3_), 20% acetonitrile (MeCN) and were disrupted by sonication (3x 10 sec sonication bursts at 62% amplitude (or 182 W power) with 10s pause between each sonication and tubes placed in ice for a 1 min pause. This was repeated three times. Tissue pieces were lysed similarly and were disrupted by sonication (10x 10 s sonication bursts as above). This was repeated three times. Proteins in cell lysates were reduced with 5 mM final concentration tris(2-carboxyethyl) phosphine at 37°C for 45 min and alkylated with a 10 mM final concentration of iodoacetamide for 45 min in dark at room temperature followed by Qubit protein quantitation to estimate protein content.

### N- and O-glycan release

For hiPSC-CM, enriched CM, and primary tissue homogenate, 64 μg of protein was immobilized on a polyvinylidene difluoride (PVDF) membrane with subsequent *N-* and *O-*glycan release as previously described [26]. For the cell culture media analysis, a maximum of 30 μL of sample was used with the aim of immobilizing 40 μg of protein. In brief, protein spots were stained with Direct Blue (Sigma-Aldrich, MO, USA). Membrane spots were excised and washed in separate wells in a flat bottom polypropylene 96-well plate. *N-*glycans were released from the membrane-bound protein using 2 U PNGase F (Promega, WI, USA) with overnight incubation at 37°C. Following *N-*glycan removal, 500 mM NaBH_4_ in a 50 mM KOH solution was added to the membrane spots for 16 hr to release reduced *O-*linked glycans by reductive β-elimination. Released *N-*glycans were reduced with 1 M NaBH_4_ in a 50 mM KOH solution for 3 hr at 50°C, after which the reaction was neutralized by adding equimolar glacial acetic acid. Both *N-*glycans and *O-*glycans were desalted and enriched offline using Dowex 50WX8 (200-400 mesh) strong cation exchange resin followed by PGC solid phase extraction micro-columns (Thermo Fisher Scientific) prior to analysis.

### N- and O-glycan data acquisition

Samples were dissolved in 65 μL 10mM NH_4_HCO_3_ containing 1 μL of dextran ladder internal standard (ISTD, 26 ng), centrifuged to remove particulates, and 60 μL was transferred to autosampler vials. For hiPSC-CM time course, a pooled QC sample was generated by combining 3 μL from each sample. Samples were batched by replicate and samples within each batch were randomized prior to injection. An internal standard only sample was analyzed between each replicate block to verify no carryover. PGC-LC-ESI-MS/MS experiments were performed using a nanoLC-2D high performance liquid chromatography system (Eksigent, CA, USA) interfaced with an LTQ Orbitrap Velos hybrid mass spectrometer (Thermo Fisher Scientific). Released glycans were separated on a PGC-LC column (3 μm, 100 mm × 0.18 mm, Hypercarb, Thermo Fisher Scientific) maintained at 80°C for *N*-glycans and 40°C for *O*-glycans (Details provided in Table S2). To enhance ionization and improve detection of low abundance glycans, post-column make-up flow consisting of 100% methanol was utilized. All methodological details are provided in Supplemental Methods.

### Glycan data analysis and library construction

Normalized retention time (RT) by dextran ladder was determined as described previously using Skyline [27] (length 4-9 for *N-*glycans and length 3-9 for *O-*glycans) and proprietary raw files were converted to an open and vendor neutral file format, mzML, using Proteowizard v3.0.8725 [28]. The pooled QC samples were used as representative files for *N-* and *O-*glycan library construction as these samples contained all glycans from one replicate sample for each timepoint. MS2 scans, required for confirmation of glycan identity, were filtered to include only precursor masses consistent with probable human glycan compositions (searched within 20 ppm error with GlycoMod [29]). For *N-*glycans, MS2 scans with assigned glucose unit (GU) values less than 3.5 GU and greater than 18 GU were removed. For *O-*glycans, MS2 scans with assigned GU values less than 2 GU and greater than 18 GU were removed. Redundant MS2 scans for each glycan structure were removed, leaving the most intense MS2 scan for each observed glycan structure. Based on the accurate precursor mass and most probable composition, chemical formulae were assigned to each MS2 scan. The MS2 scan metadata were then used to develop a transition list for Skyline-based quantitation.

### Glycan quantitation

Skyline v4.2.1.19095 [30] was used for all glycan quantitation. 99.999% of the isotopic envelope with a centroid mass accuracy value of 20 ppm was used for peak integration for glycan structure quantitation. All observed charge states for each glycan were used for quantitation. Peak picking was largely automated based on explicit retention time and integration bounds were manually supervised. Peak areas for each quantified glycan structure were exported and normalized to relative signal in Excel. Glycan structure relative signal was calculated as the specific glycan structure’s peak area as a percentage of the total peak area for all glycan structures quantified for each sample.

### Glycan structure assignment

MS2 scans representing detected glycan precursors were assigned glycan structures based on the existence of A-, B-, C-, X-, Y-, and Z-product ions matched using GlycoWorkBench v2.1 [31] (available from https://code.google.com/archive/p/glycoworkbench/, maximum of 3 glycosidic cleavages and 1 cross-ring cleavage with 0.6 Da mass accuracy). For all MS/MS scans, at least two probable glycan structures were used for comparative annotation to assign the most suitable structure assignment. Diagnostic ions were used to confirm glycan motifs as described previously [32, 33]. Simplified examples of annotated MS2 spectra are represented as Figure S10.

### Glycoproteomic data analysis

An *N-*glycan composition library was constructed by combining the Byonic (v3.4.0, Protein Metrics, CA, USA) standard glycan library with a list of the glycan compositions reduced from the structures identified in this study. Raw files were downloaded from PXD005736 and searched in Byonic with the following settings: Protein database file (Human Swiss-Prot retrieved 2018), decoys used, fully tryptic peptides, no missed cleavages, 10 ppm precursor mass tolerance, 20 ppm product ion mass tolerance, fragmentation type HCD. Fixed modification: Carbamidomethyl on C. Rare (variable) modifications: Acetyl (N-term), Oxidation (M), Deamidation (N) and N-glycosylation (NXS/T). Glycopeptide spectral matches identified by Byonic were manually inspected for confirmatory product ions (Y1 and oxonium ions [34]).

#### Immunohistochemistry

Cryosections of a right ventricle heart section were prepared by the Children’s Research Institute Histology Core. Concanavalin A conjugated to a fluorophore, Cy3 (Vector Laboratories, Burlingame, California) was diluted with Tyrode solution to a 50 µg/mL concentration. Sections to be labelled with ConA were washed with Tyrode solution, then labelled with ConA-Cy3 (15 µg per section) for 1 hr at room temperature followed by Hoechst staining (500 µg per section) for 10 min. Sections to be labelled with Wheat Germ Agglutinin (WGA) were washed with PBS, then labelled with WGA-Alexa 568 (150 µg per section) for 1 hr at room temperature followed by Hoechst staining (500 µg per section) for 10 min. Sections were imaged with Nikon Elements-D 3.1 software using a Nikon A1R confocal microscope.

### Data Availability

To promote utility of the glycan structure libraries generated here and adhere to MIRAGE [35, 36] guidelines, all raw and processed data, annotated MS/MS spectra and instrument metadata are available on Glycopost (GPST000030) [37] and Panorama Public [38] (https://panoramaweb.org/GlycoCM.url). Supplementary file S1 lists all identified structures with their composition and structural features.

## Results

### An optimized analytical platform provides deep glycan structure characterization of human cardiomyocytes

Following release of *N-* and *O-*glycans from glycoproteins, structures were separated and detected using a PGC-LC-MS configuration that incorporates post-column make-up flow [39]. Advantages of this approach are that PGC enables quantification of structural isomers that cannot be resolved by MS alone and the make-up flow achieves an average of a 6-fold increase in glycan signal intensity over the standard method (Figure 2A). Demonstrating utility of this approach, 95 *N*-glycan structures, which represent all major *N-*glycan biosynthetic classes over 3 orders of magnitude (Figure 2B), and 4 *O*-glycan structures (Figure S3) were identified in primary enriched CM and heart tissue. Altogether, there was no clear relationship between glycan class and relative signal as structures from the high mannose and complex bi-antennary classes spanned 3 orders of magnitude; however, paucimannosidic and complex mono-antennary class structures both had relative signals less than 1%. Based on signal-to-noise ratios, it is predicted that 42 glycan structures would have been undetectable without the use of the optimized post-column make-up flow configuration (Figure 2B, Figure S0).

**Figure 2.**
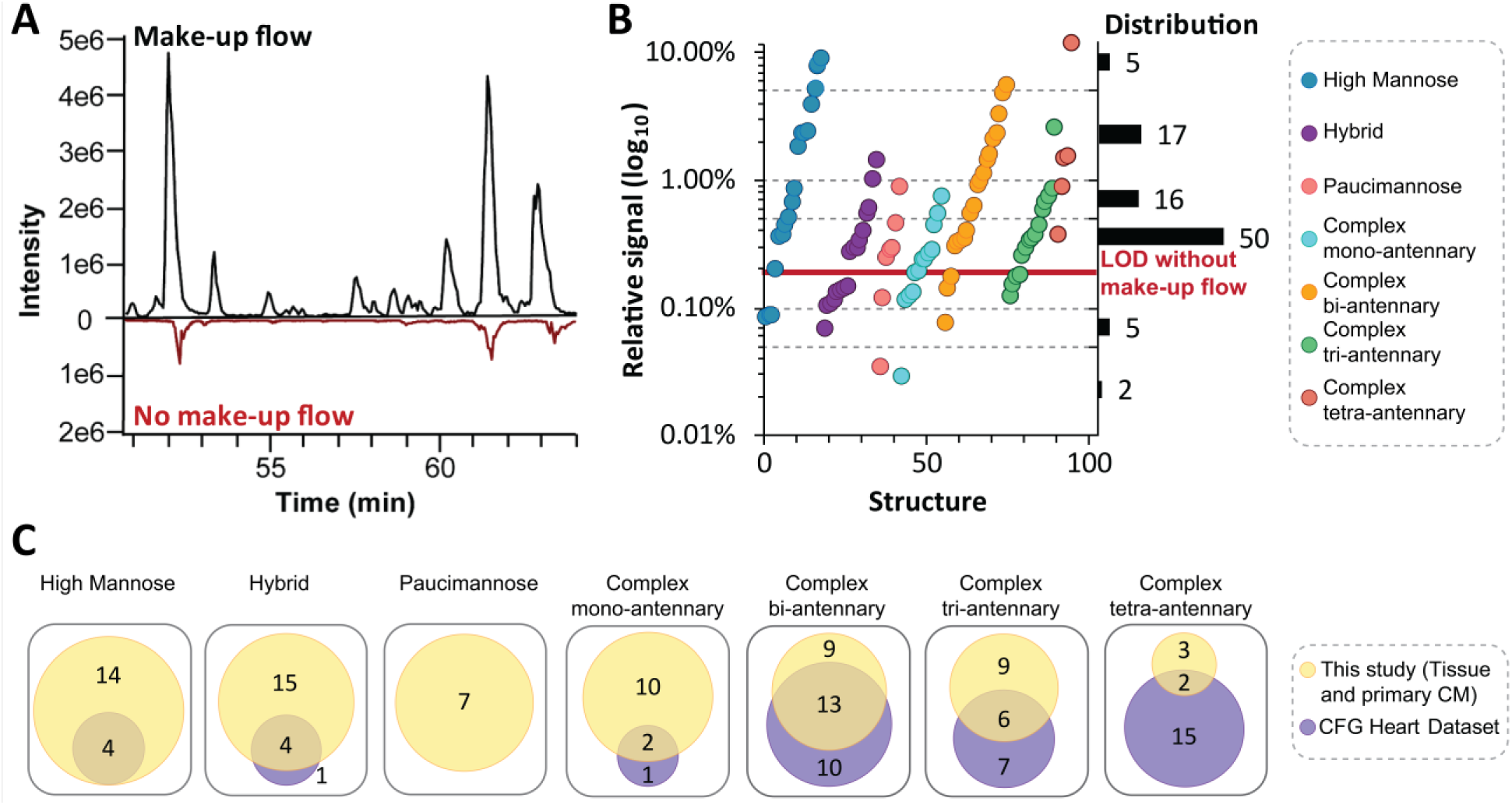
Optimization of LC-MS parameters expand the dynamic range for improved detection of glycan structures. **A** Base peak chromatogram of separated glycan structures with and without make-up flow. **B** Scatter plot of average glycan structure signal for each biosynthetic class from heart tissue and primary enriched CM with distribution shown on right. **C** Number of quantified glycan structures across biosynthetic classes for heart tissue and primary CM for our study in reference to CFG Heart Dataset [21].

Having established that our method provides a sensitive view of the glycome, we assessed the orthogonality of our data to those from a previous report by the Consortium of Functional Glycomics (CFG) [21], which applied matrix assisted laser desorption/ionization tandem mass spectrometry (MALDI-MS/MS) to identify 104 derivatized *N-*glycan compositions from >100 mg of homogenized human heart tissue. Overall, our analysis of heart tissue and CM enriched from primary heart tissue identified 98 *N*-glycan structures which covered the majority of *N-*glycans identified by the CFG (Figure 2C, 64%). The high mannose, hybrid and complex mono-antennary classes, found in the early stages of the protein glycosylation biosynthetic pathway, had over 90% overlap between studies; however, the paucimannose class, a newly validated glycan class in humans [40], was only detected in our study. In contrast, only 25% of the complex tetra-antennary class reported by the CFG were detected and characterized in our study. The differences in glycans identified between these studies could be due to differences in amount of starting material (64 µg vs >100 mg), differences in analyte (derivatized vs native) and ionization technique (ESI vs MALDI) [41]. Another possible confounding difference is patient blood groups which result in unique glycan structures [42].

### Quantitative differences between primary enriched CMs and heart tissue homogenate

To identify glycan structures that can be specifically attributed to CM, we compared the *N*-glycome of CM enriched from primary heart tissue to that of tissue homogenate from adjacent sections of the same donor hearts. Of the quantified glycan structures, 21 (22%) were significantly different (p < 0.05 and greater than 2-fold difference in CM/tissue average signal, Figure 3A) between enriched CM and homogenized heart tissue. Sixteen glycans were found with higher signal in the enriched CM, including 12 structures from the high mannose class (ranging from 2- to 6-fold differences; Figure 3B). Of the five glycans with higher signal in the tissue homogenate, three structures feature the LacDiNAc motif (2- to 7-fold differences; Figure 3C). Notably, the LacDiNAc motif is difficult to elucidate with a composition-based approach [43] and the identification of these structures based on their unique fragmentation pattern highlights an advantage of our structure-based approach. The high mannose *N*-glycans which correlated to enriched CM represent the beginning of the glycan biosynthetic pathway and can be found attached to glycoproteins transitioning through the glycosylation pathway and glycoproteins with insufficient solvent accessibility for further biosynthetic processing [44]. We verified these differences are not artifacts of residual enzymatic activity that is occurring during sample processing but are a result of cell-type enrichment (Figure S2). In contrast, the glycan structures enriched in tissue homogenate (*i.e.* prevalent in non-CM cell types) have undergone further processing involving the addition of GalNAc to the glycan non-reducing terminus by β4GalNAc-T3 [45].

**Figure 3.**
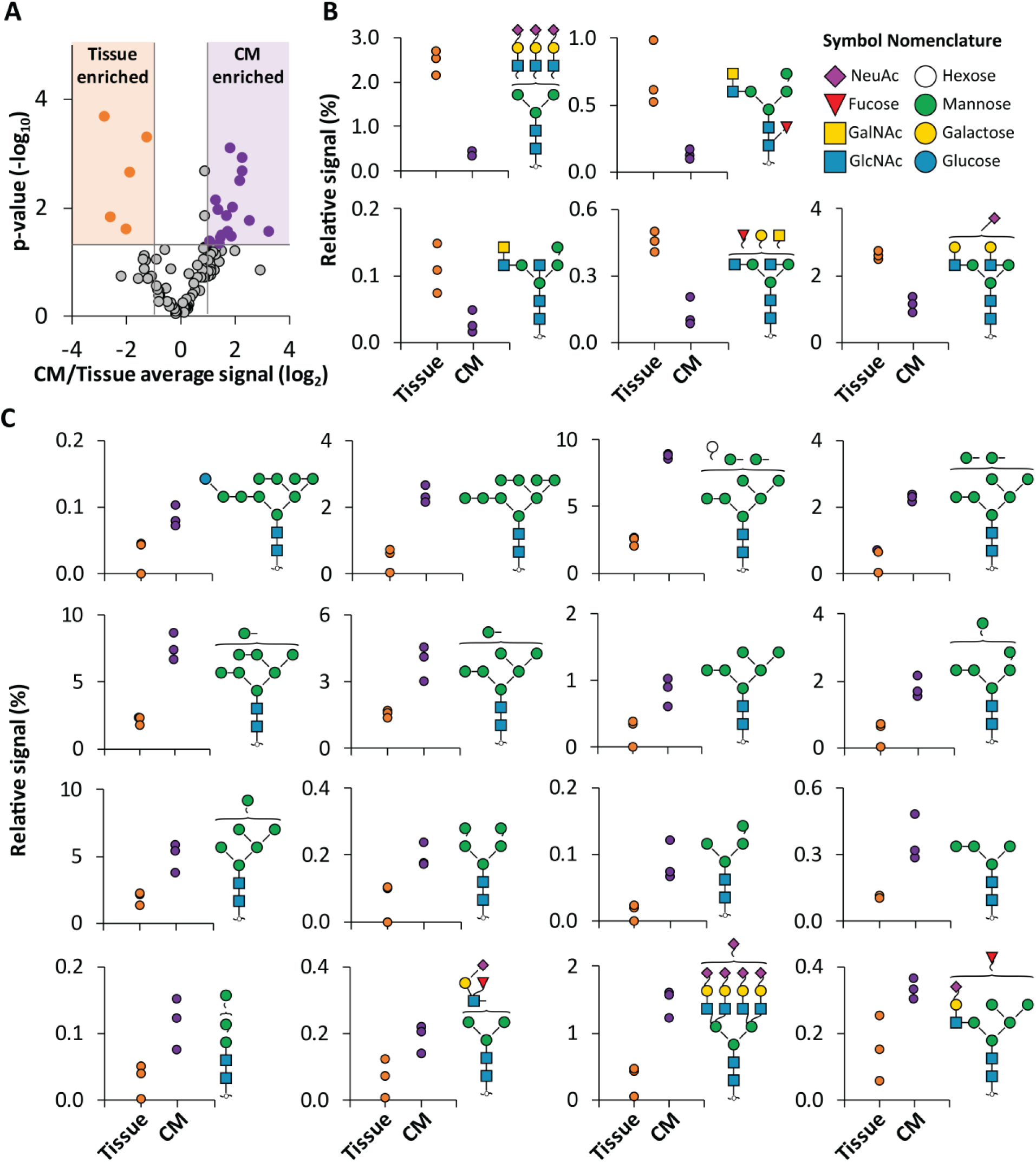
Comparison of *N*-glycans in primary enriched CM and tissue homogenate. **A** Volcano plot of quantified *N*-glycans with thresholds (shading) applied for significance (p<0.05) and fold change (>2 fold) between heart tissue homogenate (n=3) and primary enriched CM samples (donor matched, n=3). Colored data points are glycans selected for display in B. **B** *N*-glycan structures that are significantly increased in the tissue homogenate. **C** *N*-glycan structures that are significantly increased in the enriched CM.

Altogether, these results demonstrate there are significant glycosylation differences among cell types within the heart, with CM preferentially exhibiting glycans that are less mature. To our knowledge, this is the first cell-type specific view of the human cardiac glycome. These data inform future studies that use genomic, transcriptomic, metabolomic, and proteomic approaches to study the genes (enzymes) and metabolites involved in protein glycosylation that are necessary for normal heart development and which are aberrant in disease by defining the specific glycan structures that are similar and different among cell types in the heart.

### Contrasting primary enriched CM and hiPSC-CM glycan structures

The use of hiPSC-CM as a model system has shown promise for a variety of applications including cardiotoxicity screening and channelopathy modeling [46, 47]. However, differences in morphology, metabolic and functional properties, transcriptome, and proteome between hiPSC-CM and primary CM have been widely reported (reviewed in [48–50]). As glycans are important for cell proliferation [51], histocompatibility of transplanted tissue [52], and maintenance of cardiac homeostasis [10, 53], *N-*glycans observed in primary enriched CM and hiPSC-CM were compared with the expectation that differences could be exploited for future cell and tissue engineering efforts to enhance the accuracy of the culture model. In these analyses, we considered data at both the structural (identity, order, and linkages of monosaccharides in the glycan) and compositional level (identity of monosaccharides in the glycan). This was included because although the structural data are expected to provide detail not revealed by a compositional view, compositional information is more facile to obtain and is achievable by a wider range of analytical techniques (*e.g.* HILIC LC-FLR/MS or CE-FLR/MS[54] on released glycans or C18 LC-MS on intact glycopeptides [55]). Therefore, we aimed to evaluate whether the benefits provided by a structure-based approach balance the additional effort necessary to achieve it. Finally, to confirm that cell culture components are not contributing to observed differences between hiPSC-CM and primary enriched CM, all culture components were subjected to glycan analysis and glycan structures specific for the culture components were identified. Subsequently, targeted data analysis confirmed these glycan structures were not found in hiPSC-CM glycan profiles (Figure S4).

Overall, 23% of the structures were shared between primary enriched CM and hiPSC-CM (Figure 4A). Sample-specific glycans are observed in every biosynthetic class, and most of the differences appear to be proportional to the size of the glycan classes, with the complex bi-/tri-/tetra-antennary classes featuring the largest number of differences. The differences in *N-*glycan profile between these samples could be related to the relative immaturity of the hiPSC-CM or the possibility that hiPSC-CM phenotype is “ventricular-like” and not specifically right ventricle as the enriched CM, among other possible differences. Nevertheless, by linking these structural data to the specific metabolites (nucleotide sugars), glycosyltransferases, and glycosidases that are required to generate these structures (Figure S9), we identify candidate enzymes responsible for the sample-specific unique structures (Figure 4B). For example, different sialylation linkages are found in primary CM compared to those structures found in the hiPSC-CM (Figure 4B). Based on the known actions of enzymes required to achieve these structures, these data reveal that a core-fucose glycosyltransferase (FUT8) and α2,6 sialic acid glycosyltransferases (ST6GAL1-2) are required for primary enriched CM, and B1,4 N-acetyl glucosamine glycosyltransferases (MGAT4A/B) is required for hiPSC-CM. These present pathways that could be targeted in future efforts aimed at generating hiPSC-CM that more closely resemble the adult phenotype. Moreover, these data illustrate why a structure-based approach is important when evaluating cell-type specific differences. Specifically, if only glycan compositions (agnostic of linkages) were considered, the diversity of glycan structures would be underestimated. For example, the complexity caused by different sialic acid linkages results in 62 unique complex tri-antennary structures but only 20 unique compositions for hiPSC-CM (Figure 4A). Overall, our data reveal 237 *N*-glycan structures corresponding to 119 *N*-glycan compositions in hiPSC-CM, including nearly all 49 *N-*glycan structures previously described for hiPSC-CM that were detected using 2D HPLC mapping [56]. Altogether, our study identifies additional structures compared to this previous view, and, among others, adds 39 complex tetra-antennary structures not previously reported in hPSC-CM.

**Figure 4.**
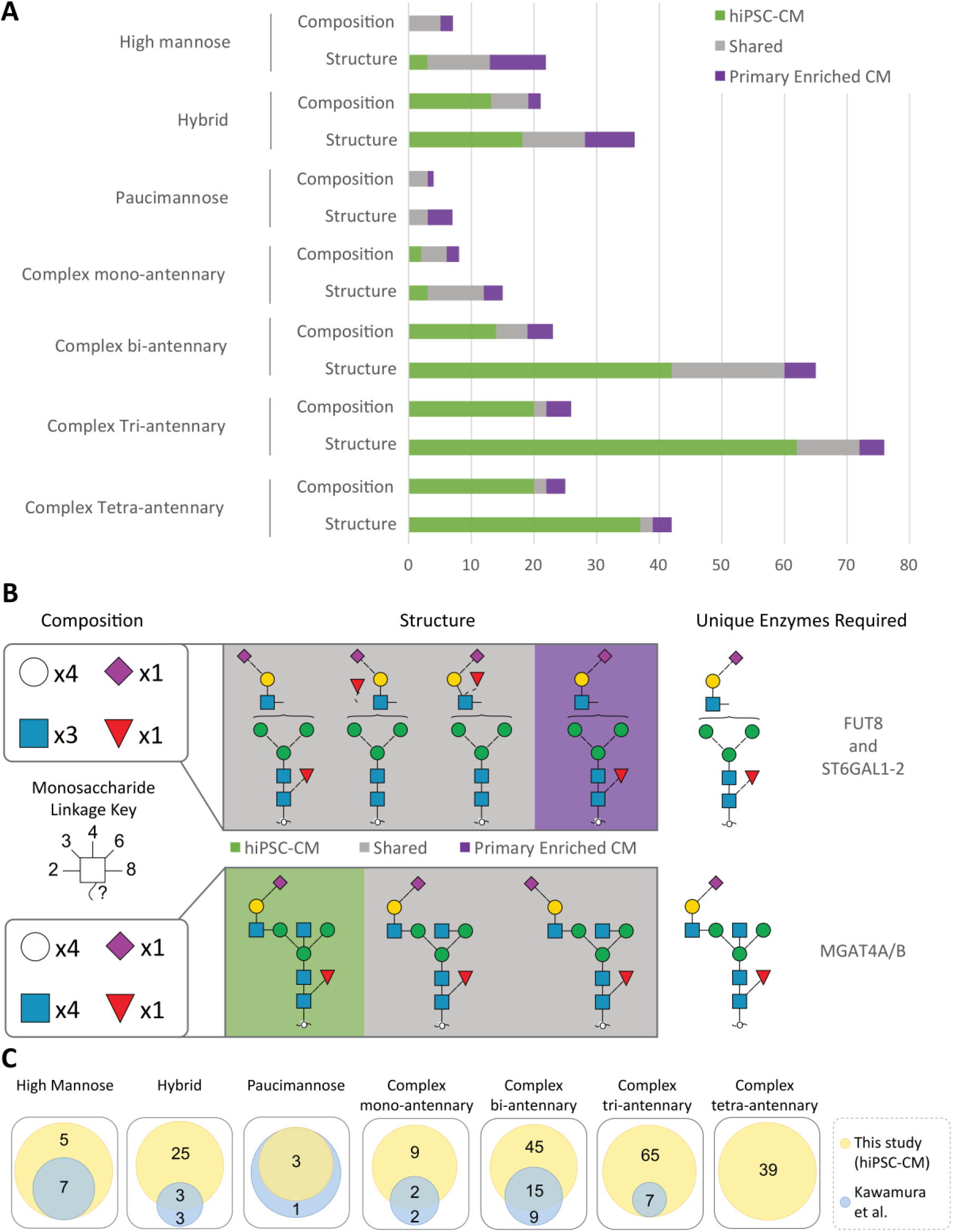
Comparison of *N-*glycan structures and classes identified in hiPSC-CM and primary enriched CM sample sets. **A** Characterized glycan structures for each sample and respective classes compared to their respective reduced compositions. **B** Examples of sample-specific and non-specific glycan structural isomers and responsible glycosyltransferases. **C** Glycan structures identified in hiPSC-CM in reference to Kawamura *et al* [56].

### Glycome dynamics during hiPSC-CM time in culture and relationship to primary CM

It is well-established that hiPSC-CM cultures are not static. Rather, their metabolic requirements and functional properties are dynamic throughout early differentiation and with extended culturing [57–60]. To complement the qualitative glycome differences between primary enriched CM and hiPSC-CM observed here and previous efforts to map the transcriptome and proteome throughout hiPSC-CM differentiation [61–64], we quantified the *N-* and *O-*glycome of hiPSC-CM throughout 20-100 days in culture. These timepoints represent cells that have undergone CM commitment and avoid changes attributed to early stages of differentiation (*e.g.* mesoderm commitment) which could confound interpretations related specifically to CM dynamics. Overall, >30% of the *N-*glycome (184 structures) significantly differs (ANOVA, p <0.01) across these timepoints (Figure 5A, Figures S6, S7) and conversely, the *O*-glycan profile does not significantly change (Figure S5). The quantitative differences in *N*-glycan structures range from 4-fold decreases to 100-fold increases (Figure 5B). The high-mannose and hybrid classes had few significant changes across the time-course, whereas all complex mono-antennary class structures that significantly changed, decreased over time. The complex bi-, tri- and tetra-antennary classes all contained structures which significantly changed over the time-course. While most structures of the bi-antennary class increased over time, the tetra-antennary class largely decreased with extended culturing. However, the directionality is not universally observed across all members of each class. For example, the complex tri-antennary class contains structures that exhibit the largest increases (e.g. fucosylated and tetra-sialylated) as well as the largest decreases (e.g. fucosylated and di-sialylated).

**Figure 5.**
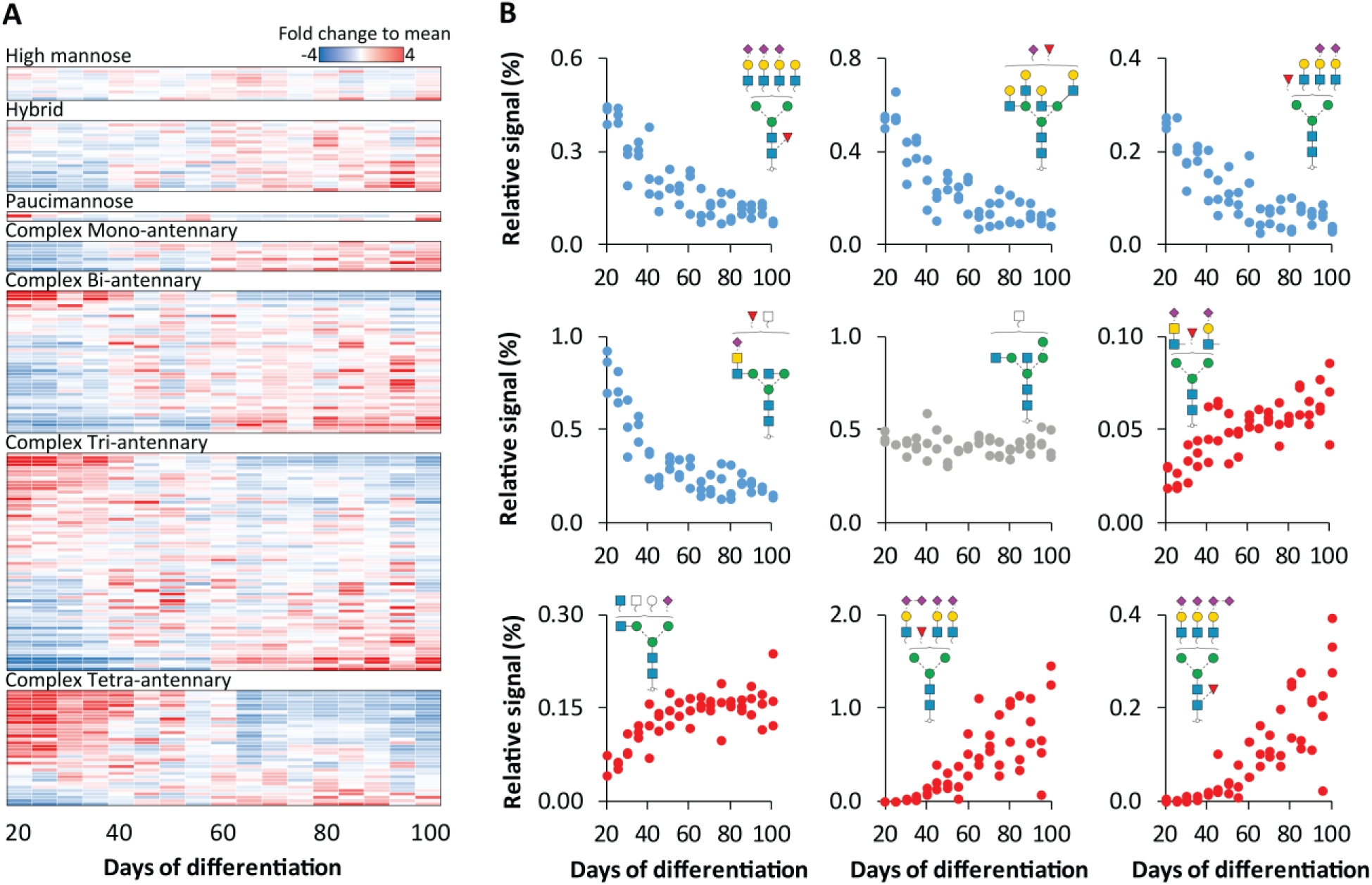
Dynamic *N*-glycome changes during hiPSC-CM differentiation. **A** Heatmap of average *N*-glycan abundance across 80 days of differentiation (n=3 for each time-point), normalized to each individual structure and classified by glycan type. **B** Scatter plot of relative signal for selected glycan structures.

To link these results to the glycan biosynthetic pathway, structural motifs (*i.e.* pattern) were evaluated. Overall, the relative signal of total sialylation, sialic acid linkages, bisecting GlcNAc and LacDiNAc structural motifs did not significantly change (Figure S8). While core-fucosylation and outer-arm fucosylation also did not significantly change, the relative motif signal for structures featuring both forms of fucosylation decreased 7-fold over the time-course (Figure S8). Notably, this observation would not be revealed by lectin arrays, as they can detect either core-fucose or outer arm fucose, but not both localized to the same structure. Altogether, these results demonstrate that the glycome is dynamic during differentiation. Considering these data in conjunction with a recent transcriptome analysis of hiPSC-CM over 90 days of differentiation, our observation that α2,8 sialylation motif increases 100-fold over time of differentiation is consistent with the increase in ST8SIA5 transcript reported previously [65]. As transcripts for the other glycosyltransferases necessary for the α2,8 sialic acid do not significantly change over time, it is possible that ST8SIA5 enzyme is specifically responsible for the increase in this motif over time, although future studies are required to confirm this. Given the biological relevance of glycosylation in cellular function and physiology, these data suggest that future studies aimed at evaluating hiPSC-CM models should consider the glycome as an important complement to other -omic data. Finally, when these dynamic changes during hiPSC-CM culture are considered in light of previous findings that protein glycosylation modulates adult *vs* neonatal CM electrophysiology [66] and there are significant protein glycosylation differences between neonatal and adult rat heart tissue [8], they support the possibility that glycan signatures may be useful for evaluating hPSC-CM maturation stage and specific structures may be necessary to achieve more mature cellular phenotypes.

To explore whether extended culturing results in a hiPSC-CM glycome that more closely resembles the adult glycan phenotype, a principal component analysis (PCA) was performed [67]. As shown in Figure 6A, hiPSC-CM across all time-points examined here are similar to each other, though increasing time in culture results in a subtle shift towards primary human CMs. As expected based on differences summarized in Figure 3, the tissue homogenate and primary enriched CM cluster by sample type rather than by patient (Figure 6A), although biological variation appears to have an impact on *N*-glycome variation. To understand the glycan structures responsible for this separation, the glycan structures that have the greatest contribution to each sample set in the PCA are annotated in a loading plot (Figure 6B). High mannose glycan structures were found to be predominantly responsible for the separation of the primary enriched CM from tissue homogenate, which is consistent with our earlier stated findings (Figure 3). Sialylated bi-antennary structures contribute towards the separation of tissue homogenate from the primary enriched CM and hiPSC-CM and the LacDiNAc motif contributes to the separation of hiPSC-CM from both primary sample types.

**Figure 6.**
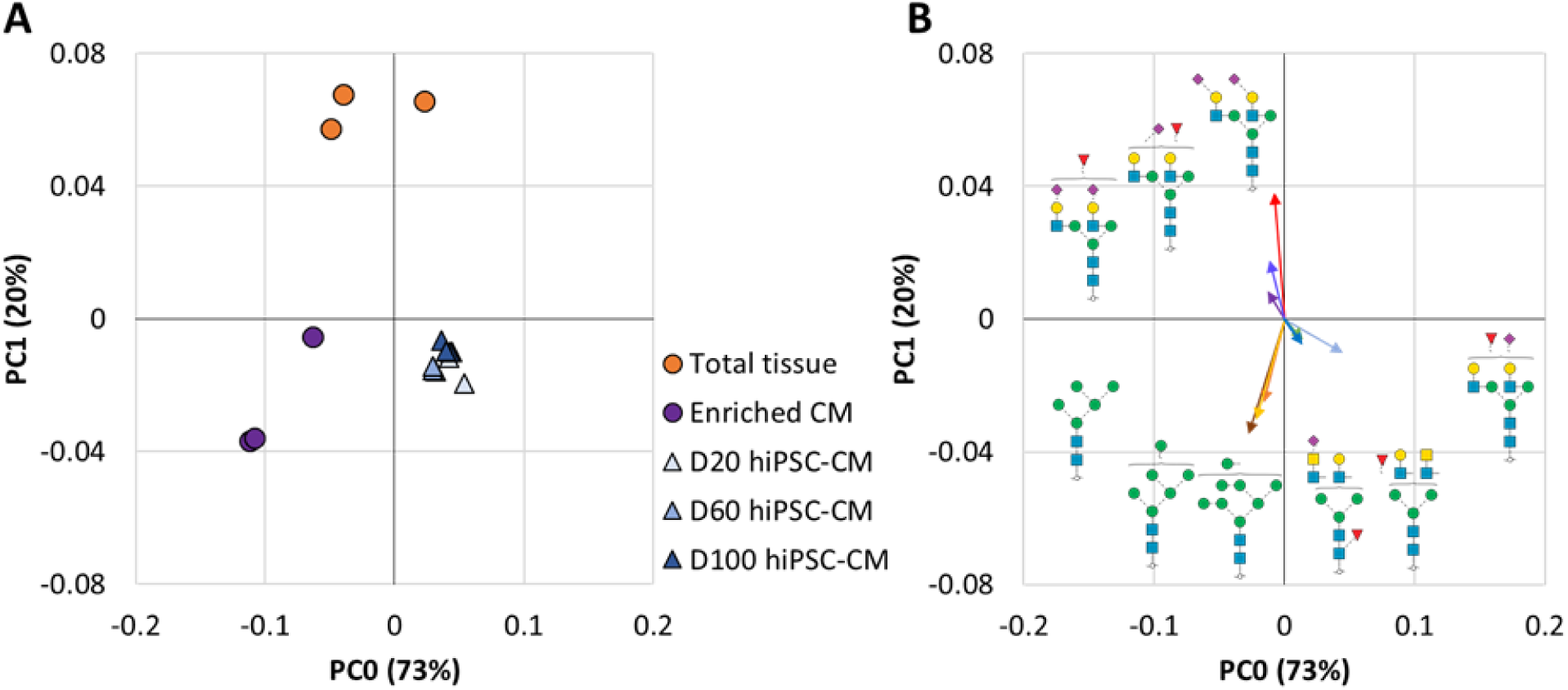
PCA analysis of *N*-glycan profiles from heart tissue homogenate (n=3), primary enriched CM (donor matched, n=3), and hiPSC-CM (n=3 for each time-point). **A** PCA plot of total *N*-glycome profiles for all sample types covering 93% of all observed variation. **B** Loading plot showing major *N*-glycan structure contributors to each cluster plotted with vectors (line length and angle) proportional to PCA dimension contribution strength and direction.

### Structural glycomic data enable orthogonal glycan-based approaches for future cardiac glycomics

Overall, our data represent the most in-depth view of the glycome of human heart tissue reported to date and provides the first glycan library that can be directly used to inform future cardiac glycoproteomic studies. By establishing a publicly available reference library of glycan structures in the human heart, including CM enriched from primary tissue, these data will support future studies of CM model comparisons and glycan-targeted immunohistochemistry (Figure 7). First, elucidation of the glycan structures and underlying compositions present on primary CM and hiPSC-CM provides a knowledge base for future efforts to identify, characterize and differentially compare glycosylated proteins present in cardiac samples. Included in this are glyco engineering efforts to increase the similarity of hiPSC-CM models to that of primary CM [68]. Second, these data can be used to inform glycoproteomic approaches aimed at characterizing proteoforms, which includes elucidating the specific glycan structures present at specific amino acid residues within a protein. Importantly, standard glycoproteomic strategies are limited to compositional information which may not distinguish among isomers [19] and glycan structure is not template derived nor predictable. Therefore, empirically derived glycan structure libraries are critical for enabling accurate glycoproteomic analyses aimed at characterizing proteoforms. To demonstrate the utility of these new cardiac glycan structure libraries for glycoproteomic applications, we analyzed the MS data from a recently described proteomic analysis of a human 3D cardiac organoid model and human heart tissue homogenate that were not originally interrogated for glycopeptides [69]. Because the MS approach used in the study is limited to informing glycan compositions, a library of 124 glycan compositions derived from this study (Figure 4) were used to inform the search space (*i.e.,* possible modifications attached to peptides).

**Figure 7.**
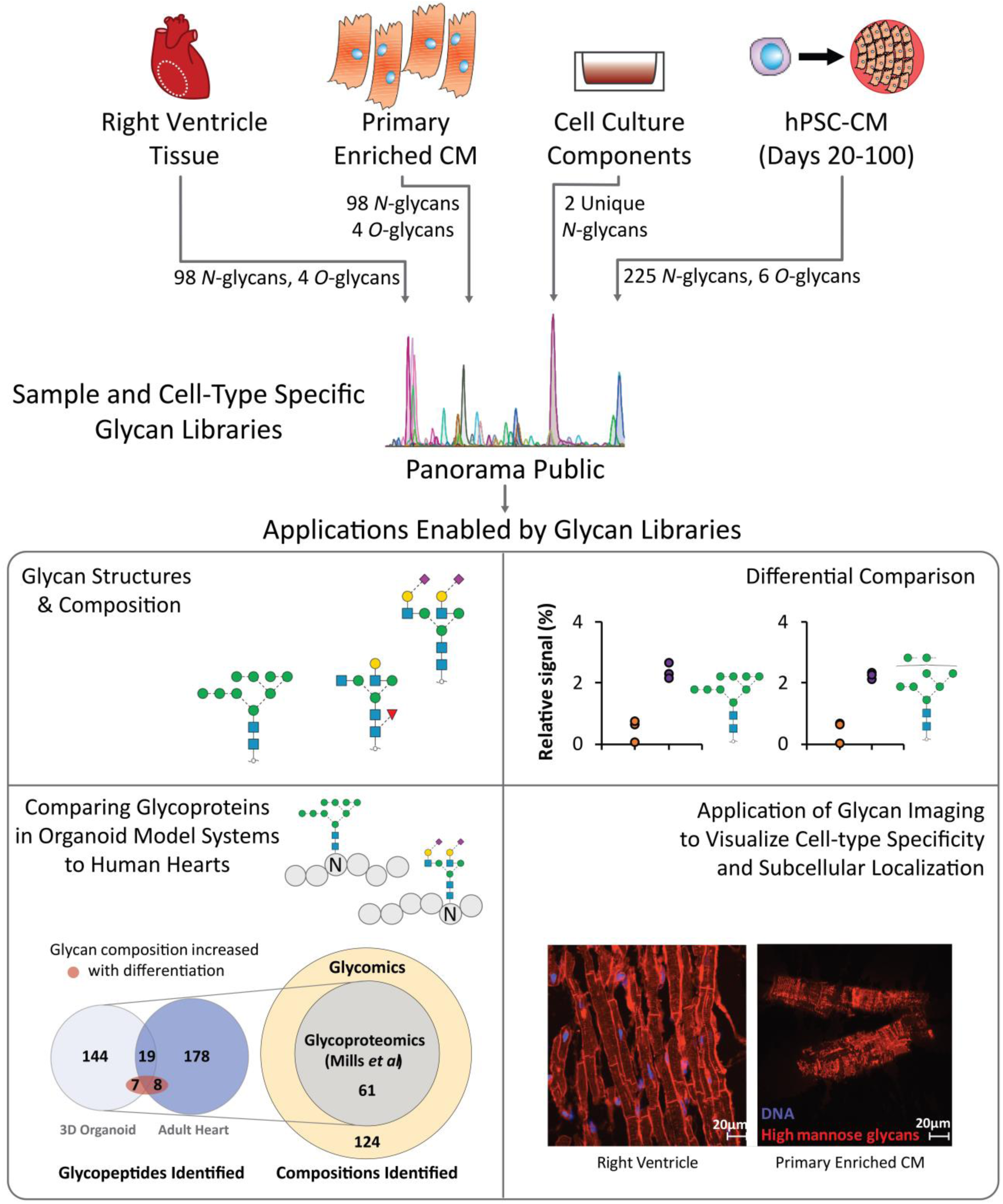
Overview of outcome and future approaches enabled by the current study. Primary CM, hiPSC-CM, and cell culture component glycan profile datasets are available in an accessible platform with future demonstrated applications towards identification of previously unidentified glycopeptides in published proteomics datasets and localization of glycan structures using lectin-based glycoimaging.

Glycopeptide spectral matches corresponding to 356 unique glycopeptides were identified. Following manual confirmatory inspection of the spectra, bona fide glycopeptides from Heparan sulfate proteoglycan 2 (HSPG2), fibronectin (FINC), and Galectin-3 binding protein (LG3BP) were detected in both the primary tissue and 3D-organoids. These proteins have been previously described as having roles in maintaining heart function and cardiovascular disease [70–72]. HSPG2 glycopeptides identified in both the heart and 3D-organoid samples feature identical glycan compositions with no detected differences in their profiles (Figure 8A). As this protein is critical for heart development and mediating injury-based vascular response [73], the conserved glycosylation between the hiPSC-CM model and the human heart may support its necessity. This contrasts with that observed for FINC and LG3BP, where different glycopeptide profiles were observed between the sample types. FINC, a protein that plays a critical role in cardiac remodeling after infarction, was found to have different glycan compositions at the same glycosite. This glycosite (N1007) has been found to be glycosylated in previous intact glycopeptide [74] and deglycosylated peptide analyses [75–77].

**Figure 8.**
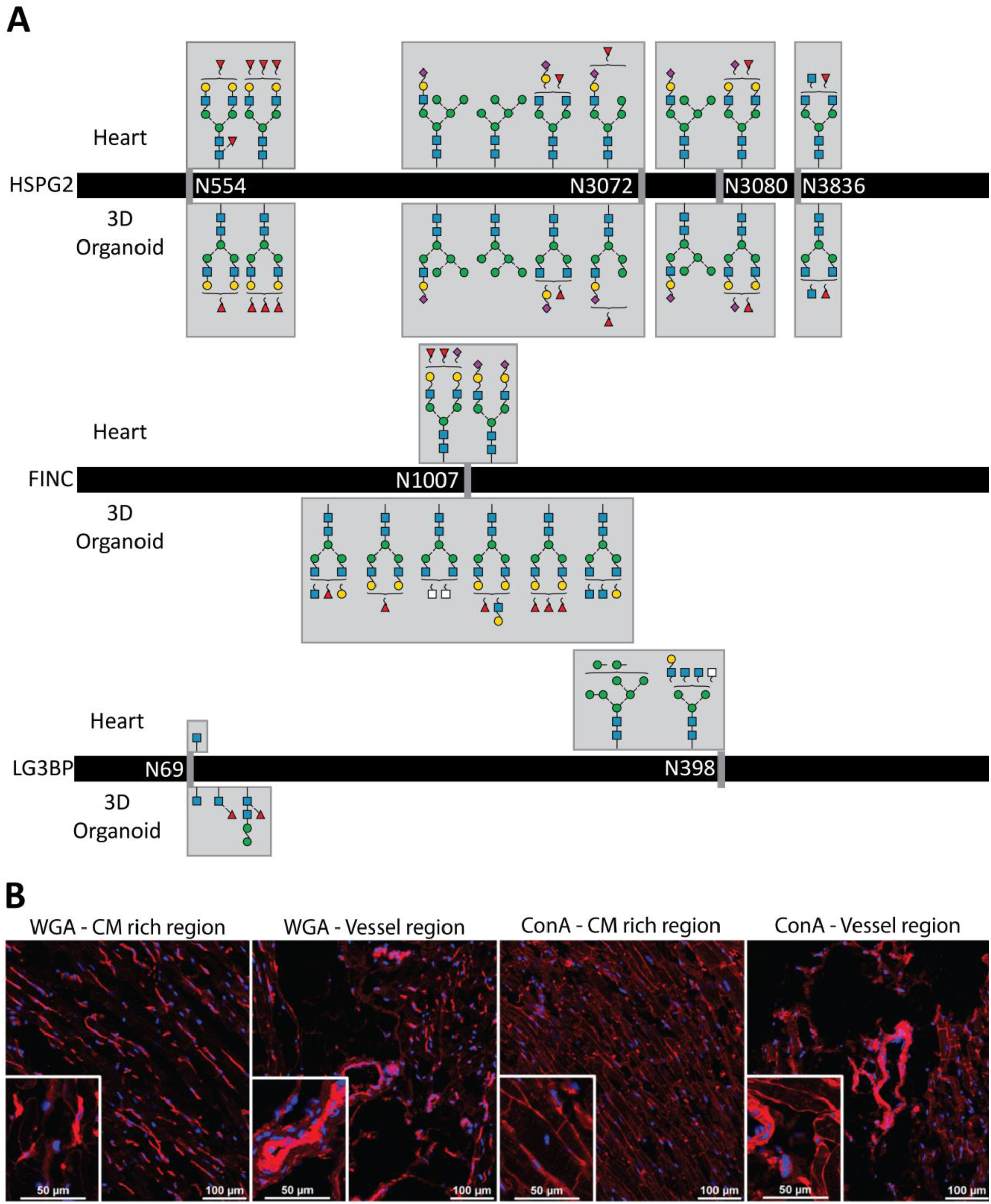
Examples of orthogonal approaches enabled by our cardiac glycan reference libraries. **A** Glycopeptide compositions identified in primary human heart vs 3D-organoid are the same for HSPG2 (top), distinct for FINC (middle) and mixed for LG3BP. **B** Immunohistochemistry images of ConA, a lectin specific for high mannose and WGA, a commonly used [80–82] membrane marker lectin specific for sialic acids.

Our results agree with those identified in the previous studies and provide structural candidates for the glycan compositions identified by intact glycopeptide analysis, including bi-antennary *N*-glycans. LG3BP, a candidate biomarker of heart failure and carotid plaques [78, 79], was found to have one glycosite with different glycan compositions between the two samples (N69) and one glycosite uniquely identified in the heart sample (N398). These glycan compositions are the first to be identified for this protein, as previous studies have only focused on deglycosylated glycopeptide analysis. Overall, our study contributes possible structural candidates for the glycoproteins discussed, catalyzing the application of these glycan structure reference libraries towards understanding the possible structural variants of glycoproteins involved in essential cardiac functions.

Beyond these data which provide further details regarding similarities and differences between primary cells and those derived *in vitro*, these glycopeptides may also be useful for interpreting the dynamic glycan profile observed here for hiPSC-CM during days 20-100 of differentiation. Specifically, 14 glycopeptides found in the Mills *et al* study [69] featured glycan compositions that were positively correlated to time of hiPSC-CM differentiation in our study (Table S4). These include HSPG2, laminin subunit alpha 2, versican core protein, and decorin – where all but two feature the concurrent core- and outer-arm fucosylation motif. The majority of these proteins are localized to the extracellular matrix, which has been demonstrated to coordinate the form and function of the developing heart [83] and regulate cardiomyocyte differentiation from pluripotent stem cells [84]. Therefore, it is possible that specific glycosylation structures on these glycoproteins are involved in, or are reflective of, cardiomyocyte maturation stage.

Finally, these glycan structure libraries will inform orthogonal methods, such as the use of lectins for array-based screening or microscopy. To demonstrate, the high mannose class found preferentially in enriched CM compared to tissue homogenate was visualized using the lectin Concanavalin A (ConA). Imaging of right ventricle heart tissue sections (Figure 7 and Figure 8B) demonstrates preferential labeling of the CM, consistent with our glycomics data. Complementing our glycomics data, imaging of tissue sections and primary enriched CM provides visualization of the high mannose structures localized to the cell surface and intracellular organelles (Figure 7), information which is difficult to discern from glycomic analyses of cell lysates. Overall, ConA is a viable alternative to a widely used lectin, WGA, which recognizes sialic acids and is widely used to visualize the plasma membrane. Whereas ConA preferentially labels CM, WGA predominantly labels non-CM and is only faintly detected on CM (Figure 8B). Altogether, ConA represents a lectin that can be used in future imaging studies to localize CM.

### Considerations for future studies

While these data present the most extensive view of the human cardiac glycome reported to date and enable the linking of identified structures to the glycan biosynthetic pathway, the analysis of released glycans does not preserve information regarding the glycoproteins to which they occupied. Therefore, future intact glycopeptide studies will be required to fully map the glycoproteome in the heart, including site-specific glycan structures in health and disease. Our method is specific for the analysis of *N-* and *O-*glycosylation and other glycosylated molecules, such as proteoglycans and glycolipids, are not detected. Finally, failure to detect specific glycan structures does not equate to their absence, rather, they could be present at levels below detection limits for this method. Regarding biological samples, the use of additional patient samples will be required to fully assess inter-patient variability, although non-diseased human heart tissue for research is a limited resource. Visual assessment of the cells obtained through our CM enrichment protocol suggest the vast majority of the cells are CM, although it remains possible that other cell types from the heart (*e.g.* endothelial cells, fibroblasts) are present in low numbers. Hence, further validation of cell type localization for glycans within tissue is required, but such efforts are currently limited by a lack of available affinity reagents for visualization. Finally, future studies using different pluripotent stem cell lines and differentiation strategies would be required to determine if the observations presented here are broadly applicable. However, having generated the glycan libraries and shared them in a public repository, future studies can use targeted analyses to rapidly and directly test hypotheses that emerge from this and other studies regarding the role of glycans in the heart and cardiomyocyte differentiation.

## Conclusions

The quantitative structural analyses of cell-type specific glycans presented here provide a key link that is necessary to understand the biosynthetic pathway of glycosylation that is present among the various cell types in the heart. Ultimately, these sample-specific glycan reference libraries provide the fundamental first step towards understanding the role glycosylation plays in cell-type specific functions and cardiac disease. The structural differences observed here, either among cell types or stages of differentiation, require complex regulation of multiple enzymes in the biosynthetic pathway, and therefore would be challenging to measure with bulk motif strategies such as lectin arrays or glycosyltransferase monitoring strategies such as RNAseq or targeted proteomics. Therefore, continued application of structure-based glycomics approaches, such as the method used here, will be essential for elucidation of the roles that glycans and glycoproteins play during developmental and disease processes in the human heart.

## Acknowledgements

This work was supported by the National Institutes of Health [R01-HL126785 and R01-HL134010 to RLG; F31-HL140914 to MW]; Funding sources were not involved in study design, data collection, interpretation, analysis or publication. Mass spectrometry analyses were performed using instrumentation in the Center for Biomedical Mass Spectrometry Research at the Medical College of Wisconsin. Tissue sectioning was performed by the Children’s Research Institute Histology core lab. Flow cytometry analyses were performed using instrumentation in the Blood Center of Wisconsin Flow Cytometry Core.

## Abbreviations

CDG: Congenital Disorders of Glycosylation
Neu5Ac: *N*-AcetylNeuraminic acid
Neu5Gc: *N*-GlycolylNeuraminic acid
MALDI: Matrix Assisted Laser Desorption Ionization
MS: Mass Spectrometry
LC: Liquid Chromatography
ESI: Electrospray ionization
PGC: Porous Graphitized Carbon
hiPSC-CM: Human induced Pluripotent Stem Cell derived Cardiomyocyte
MeCN: Acetonitrile
PVDF: PolyVinylidene DiFluoride
CFG: Consortium of Functional Glycomics
GlcNAc: *N*-Acetyl Glucosamine
LacDiNAc: *N*-Acetyl Glucosamine attached to *N*-Acetyl Galactosamine
PCA: Principal Component Analysis
WGA: Wheat Germ Agglutinin
ConA: Concanavalin A

