## Supplementary material for "Reference glycan structure libraries of primary human cardiomyocytes and pluripotent stem cell-derived cardiomyocytes reveal cell-type and culture stage-specific glycan phenotypes": S1

**Table of Contents**


| Content | Page |
|---|---|
| Supplemental Methods | S2 |
| **Figure S-0** Effect of make-up flow on glycan structure limit of detection | S5 |
| **Figure S-1** Examples of purity assessment for enriched primary CM and hiPSC-CM. | S6 |
| **Figure S-2** Cardiomyocyte enrichment artifact evaluation. | S7 |
| **Figure S-3** Identified *O*-glycan structures in primary CM and whole heart tissue | S8 |
| **Figure S-4** Targeted data analysis evaluating the presence of a culture component non-human *N*-glycan structure (di-gal motif) in hiPSC-CM samples. | S8 |
| **Figure S-5** Identified *O*-glycan structures in hiPSC-CM and relative signal of quantitative glycan structures over the differentiation period | S9 |
| **Figure S-6** Scatter plot of relative signal for *N*-glycans from hiPSC-CM significantly positively correlated (>0.7) to days of differentiation | S9 |
| **Figure S-7** Scatter plot of relative signal for *N*-glycans from hiPSC-CM significantly negatively correlated (<-0.7) to days of differentiation | S10 |
| **Figure S-8** Majority of glycan structural motifs do not majorly change for hiPSC-CM across days of differentiation. | S11 |
| **Figure S-9** Map of the enzymes and metabolites responsible for identified and quantified glycan structures of homogenized heart tissue, primary CM and hiPSC-CM | S12 |
| **Figure S-10** Example of annotated MS2 spectra for several glycan structures in Figures 4 and 5 | S14 |

### Supplemental Methods

| Table S1. Donor information | | | |
|---|---|---|---|
| **Anonymized Donor Identification Number** | **Sex** | **Age** | **Cause of Death** |
| 10960 | F | 75 | Stroke |
| 11225 | M | 58 | Gunshot wound to head |
| 11414 | M | 57 | Intracranial hemorrhage |

| Table S2. Chromatography and MS Instrument Acquisition Settings | | |
|---|---|---|
| | ***N*-glycan analysis** | ***O*-glycan analysis** |
| **Sample Amount for glycan release** | 64 µg of protein | |
| **Sample Volume Prepared, Injected** | 59 µL dried glycans dissolved in solvent A + 1 µL Dextran Ladder ISTD, 20 µL | |
| **Injection Mode** | Full Loop Direct Inject | |
| **Sample Loop** | 20 µL | |
| **Stationary Phase** | Thermo Scientific Hypercarb PGC 250 Å, 180 µm x 10 cm, 3 µm | |
| **LC Solvent A** | 100% $H_2O$<br>10 mM Ammonium Bicarbonate | |
| **LC Solvent B** | 90% MeCN,<br>10 mM Ammonium Bicarbonate | |
| **LC Gradient** | 0-25% B in 70 min<br>100% B for 10 min<br>0% B for 10 min | |
| **LC Flow Rate** | 2 µL/min | |
| **Column Temperature** | 80 ºC | 40 ºC |
| **Make-up Solvent** | 100% MeOH | |
| **Make-up Flow Rate** | 2 uL/min for 75 min<br>1 uL/min for 15 min | |
| **Mass Spectrometer** | Thermo Orbitrap Velos | |
| **Method Type** | Top 9 Data Dependent MS2 | |
| **Spray Voltage** | 2 kV | |
| **$MS^1$ Detector** | Orbitrap | |
| **$MS^1$ Scan Range** | 570-2000 *m/z* | 500-2000 *m/z* |
| **$MS^1$ Resolution** | 15,000 @ 200 m/z | |
| **$MS^1$ AGC Target** | 1e6 | |
| **$MS^1$ Maximum IT** | 100 ms | |
| **$MS^2$ Detector** | IonTrap | |
| **$MS^2$ Scan Range** | Auto Normal | |
| **$MS^2$ Resolution** | Normal (0.5 *m/z* FWHM) | |
| **Isolation Window** | 2 *m/z* | |
| **$MS^2$ AGC Target** | 1e5 | |
| **$MS^2$ Maximum IT** | 150 ms | |
| **Activation Type / Collision Energy** | CID 33%, Wideband Activation | |
| **Minimum Signal Req.** | 50 | |
| **Dynamic Exclusion** | 30 s,10 Repeats, 5 s Exclusion Duration | |

**Table S3. Flow cytometry sample preparation and data acquisition details.**

| **Sample Information** | |
|---|---|
| **Cell type(s)** | hiPSC-CM |
| **Cell line(s)** | DF6-9-9T |
| **Passage #** | |
| **Dissociation Conditions:** | 1 mL of 0.5U/mL Liberase-TH (Sigma #5401135001), 50U/mL DNase I (Sigma #10104159001) in RPMI (Thermofisher #11875-093) for 30 min at 37 °C, followed by addition of 1 mL of TrypLE (TrypLE Thermofisher #12605-010) for 5 min at 37 °C |
| **Total Cell Counts** | 6.5 – 13 x $10^6$ |
| **# of Cells per Tube** | 1 x $10^6$ |

| **Protocol Steps** | **Time** | **Reagents** | **Recipe, Catalog #s** |
|---|---|---|---|
| **1. Fixation** | 20 min | **Wash Solution** | DPBS-/- (Sigma #D8537) |
| **2. Wash** | two x 3 mL | **Fixation Solution** | 2% Formaldehyde (w/v) (ThermoFisher #28906) in 1X DPBS-/- |
| **3. Permeabilization** | Performed as one 15 min incubation | **Block Solution** | 0.5% w/v BSA (Sigma #A7906) in DPBS-/- |
| **4. Wash** | | **Permeabilization** | 0.5% Saponin (w/v) (Sigma #47036) in Block Solution |
| **5. Block** | | **Resuspension Solution** | 0.5% w/v BSA in DPBS-/- |
| **6. 1°Antibody** | 45 min | | |
| **7. Wash** | two x 3 mL | | |
| **8. Resuspension** | 500 µL | | |

**Antibodies**

| **Target** | **Clone** | **Vendor** | **Catalog #** | **Lot #** |
|---|---|---|---|---|
| Troponin T2 | 1C11 | Abcam | ab105439 | various |
| Isotype control | - | eBiosciences | 11-4714 | various |

| **Instrument Configuration** | |
|---|---|
| **Instrument** | BD LSR II |
| **Laser line** | 488nm (50mw) |
| **Emission filter** | 525/50 |
| **Fluorochrome** | FITC |

**Table S4. Glycopeptides identified from Mills et al. which feature glycan compositions that were positively correlated to time of hiPSC-CM differentiation in our study.**

| Protein | Peptide | Starting AA | Glycan Composition | Sample |
|---|---|---|---|---|
| **CERU_HUMAN** | K.EN[+1914.697]LTAPGSDSAVFFEQGTTR.I | 396 | HexNAc(4)Hex(5)Fuc(2) | **Heart tissue** |
| **HPT_HUMAN** | K.VVLHPN[+1914.697]YSQVDIGLIK.L | 236 | HexNAc(4)Hex(5)Fuc(2) | **Heart tissue** |
| **A1AT_HUMAN** | K.YLGN[+1914.697]ATAIFFLPDEGK.L | 268 | HexNAc(4)Hex(5)Fuc(2) | **Heart tissue** |
| **FETUA_HUMAN** | K.AALAAFNAQNN[+1914.697]GSNFQLEEISR.A | 166 | HexNAc(4)Hex(5)Fuc(2) | **Heart tissue** |
| **HRG_HUMAN** | R.VIDFN[+1914.697]C[+57.021]TTSSVSSALANTK.D | 121 | HexNAc(4)Hex(5)Fuc(2) | **Heart tissue** |
| **PGS2_HUMAN** | K.LGLSFNSISAVDN[+2100.761]GSLANTPHLR.E | 250 | HexNAc(5)Hex(4)Fuc(1)NeuAc(1) | **3D Organoid** |
| **CO6A2_HUMAN** | R.GTFTDC[+57.021]ALAN[+1914.697]MTEQIR.Q | 131 | HexNAc(4)Hex(5)Fuc(2) | **Heart tissue** |
| **CSPG2_HUMAN** | R.FEN[+1914.697]QTGFPPPDSR.F | 328 | HexNAc(4)Hex(5)Fuc(2) | **3D Organoid** |
| **LAMA2_HUMAN** | R.YMQN[+1914.697]LTVEQPIEVK.K | 2645 | HexNAc(4)Hex(5)Fuc(2) | **3D Organoid** |
| **LAMA2_HUMAN** | K.N[+1914.697]ESGIILLGSGGTPAPPR.R | 2558 | HexNAc(4)Hex(5)Fuc(2) | **3D Organoid** |
| **BCAM_HUMAN** | R.TQN[+1914.697]FTLLVQGSPELK.T | 437 | HexNAc(4)Hex(5)Fuc(2) | **Heart tissue** |
| **PGBM_HUMAN** | R.SLTQGSLIVGDLAPVN[+1914.697]GTSQGK.F | 3765 | HexNAc(4)Hex(5)Fuc(2) | **3D Organoid** |
| **PGBM_HUMAN** | R.NQELEDNVHISPN[+1752.645]GSIITIVGTRPSNHGTYR.C | 3060 | HexNAc(4)Hex(5)Fuc(2) | **3D Organoid** |
| **COEA1_HUMAN** | R.SFMVN[+1914.697]WTHAPGNVEK.Y | 368 | HexNAc(4)Hex(5)Fuc(2) | **Heart tissue** |

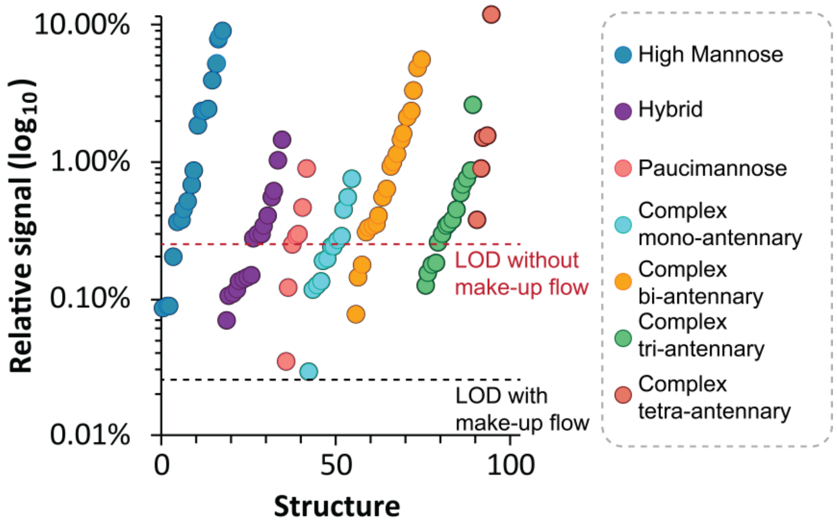


**Figure S0.** Effect of make-up flow on glycan structure limit of detection

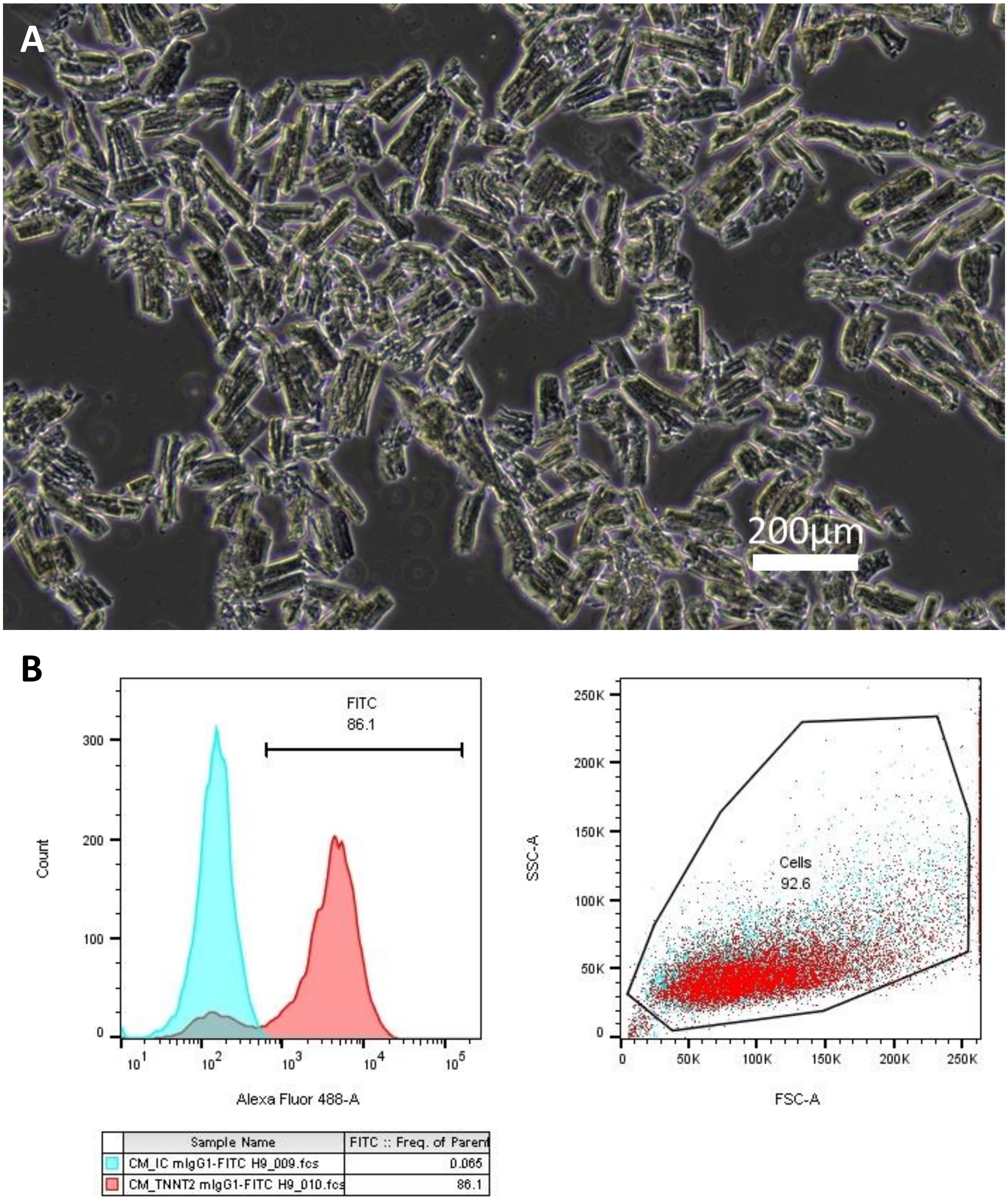


**Figure S1.** Examples of quality assessment for enriched primary CM and hiPSC-CM. **A** Bright-field image of cardiomyocyte-enriched samples from human heart tissue. **B** Representative example of percent troponin positivity in hiPSC-CM as determined by flow cytometry (isotope control is represented in blue, antibody for troponin is represented in red)

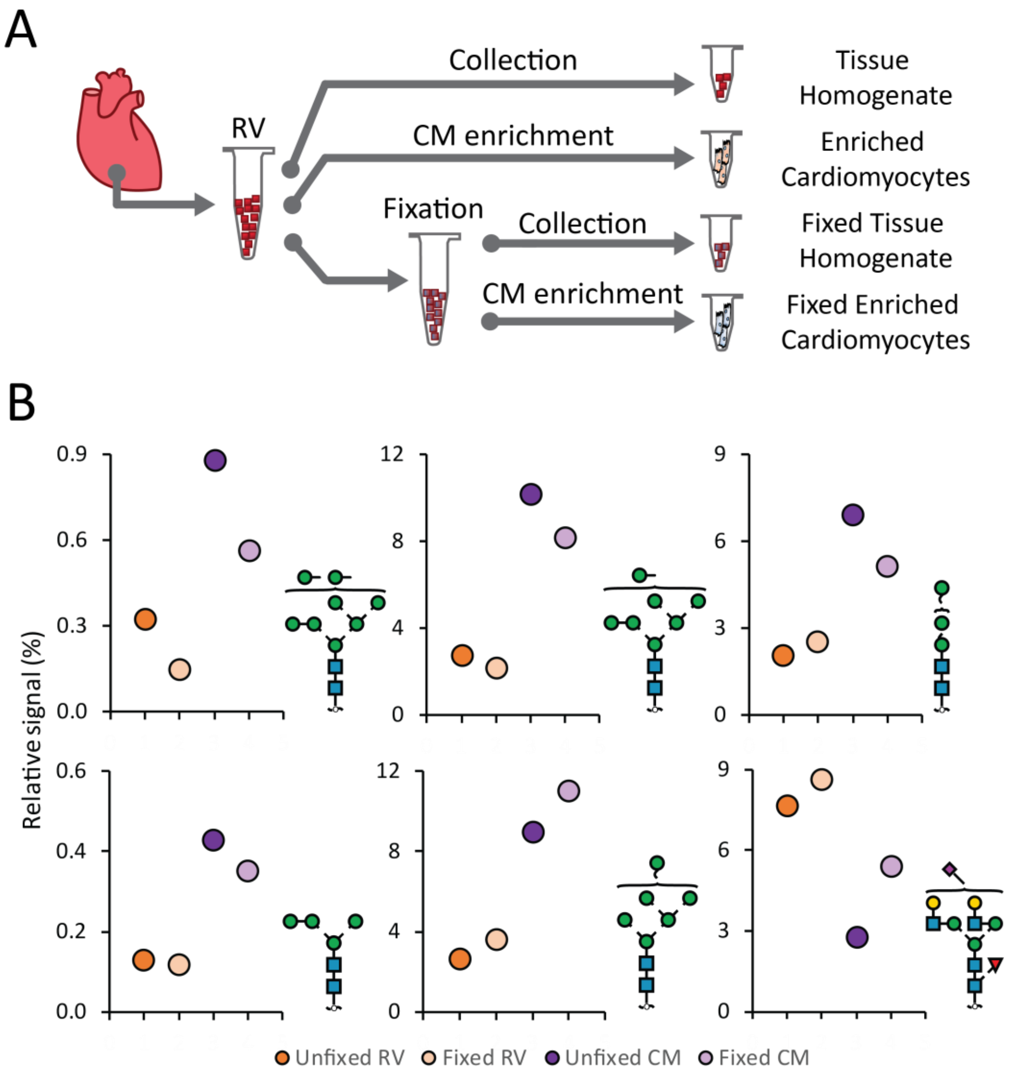


**Figure S2.** Evaluation of the possibility of artifacts owing to residual enzymatic activity during CM enrichment. **A** Experimental design to compare fixed and unfixed heart tissue homogenate and enriched CM. **B** Quantitative analysis of enriched CM glycans compared to cardiac tissue homogenate. For each sample type, no major differences are observed between unfixed and fixed samples.

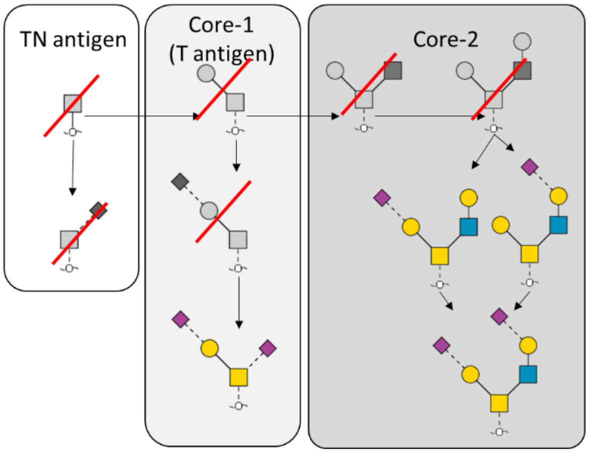


**Figure S3.** Identified *O*-glycan structures in primary CM and whole heart tissue

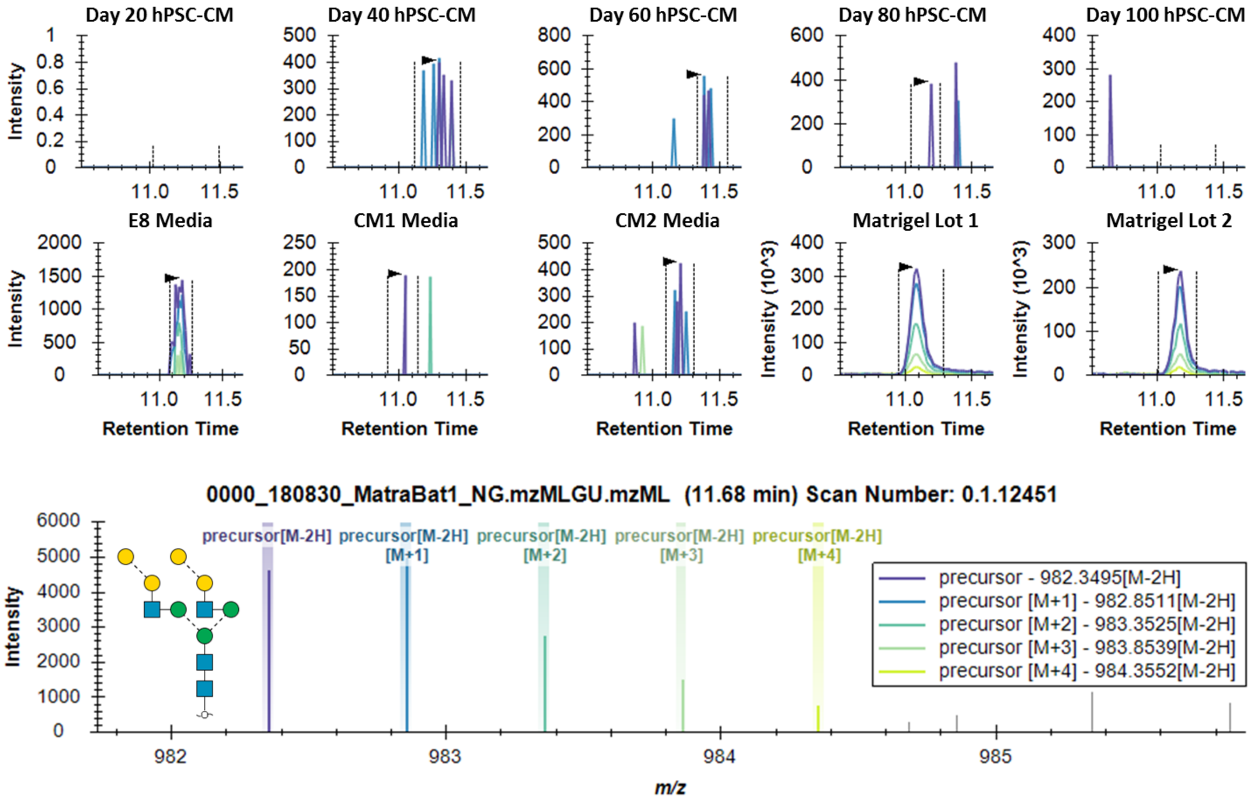


**Figure S4.** Targeted data analysis evaluating the presence of a culture component non-human *N*-glycan structure (di-gal motif) in hiPSC-CM samples

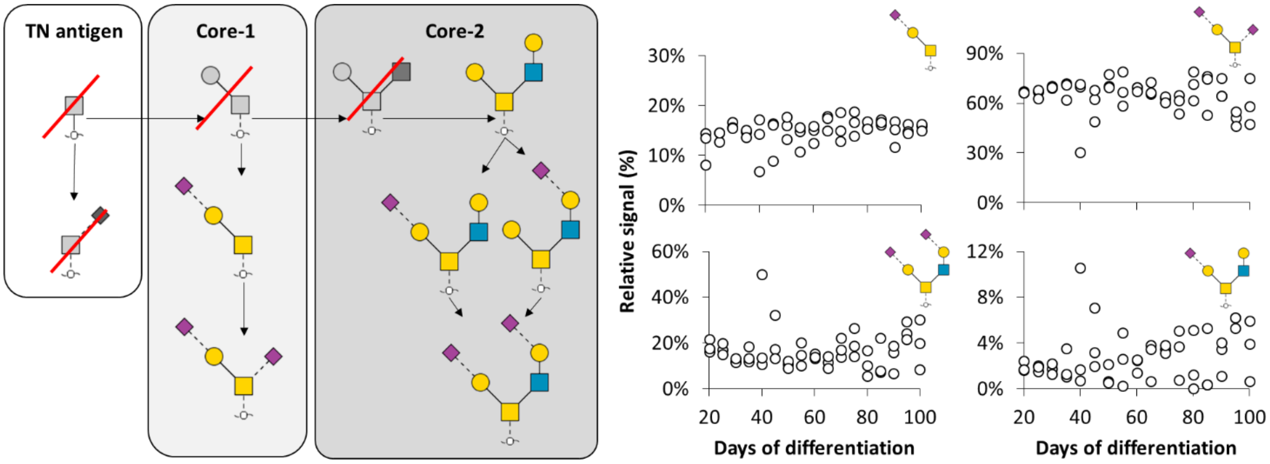


**Figure S5.** Identified *O*-glycan structures in hiPSC-CM and relative signal of quantitative glycan structures over the differentiation period

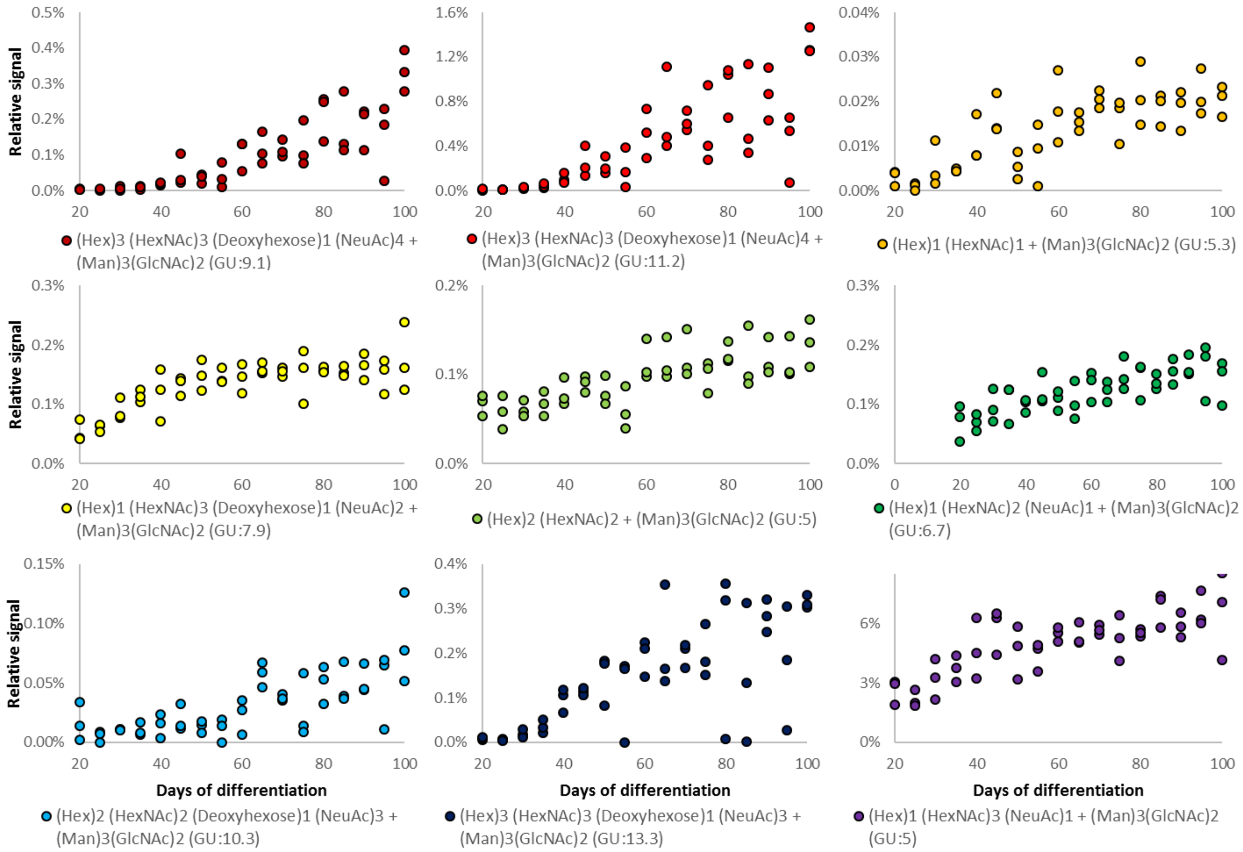


**Figure S6.** Scatter plot of relative signal for *N*-glycans from hiPSC-CM significantly positively correlated (>0.7) to days of differentiation

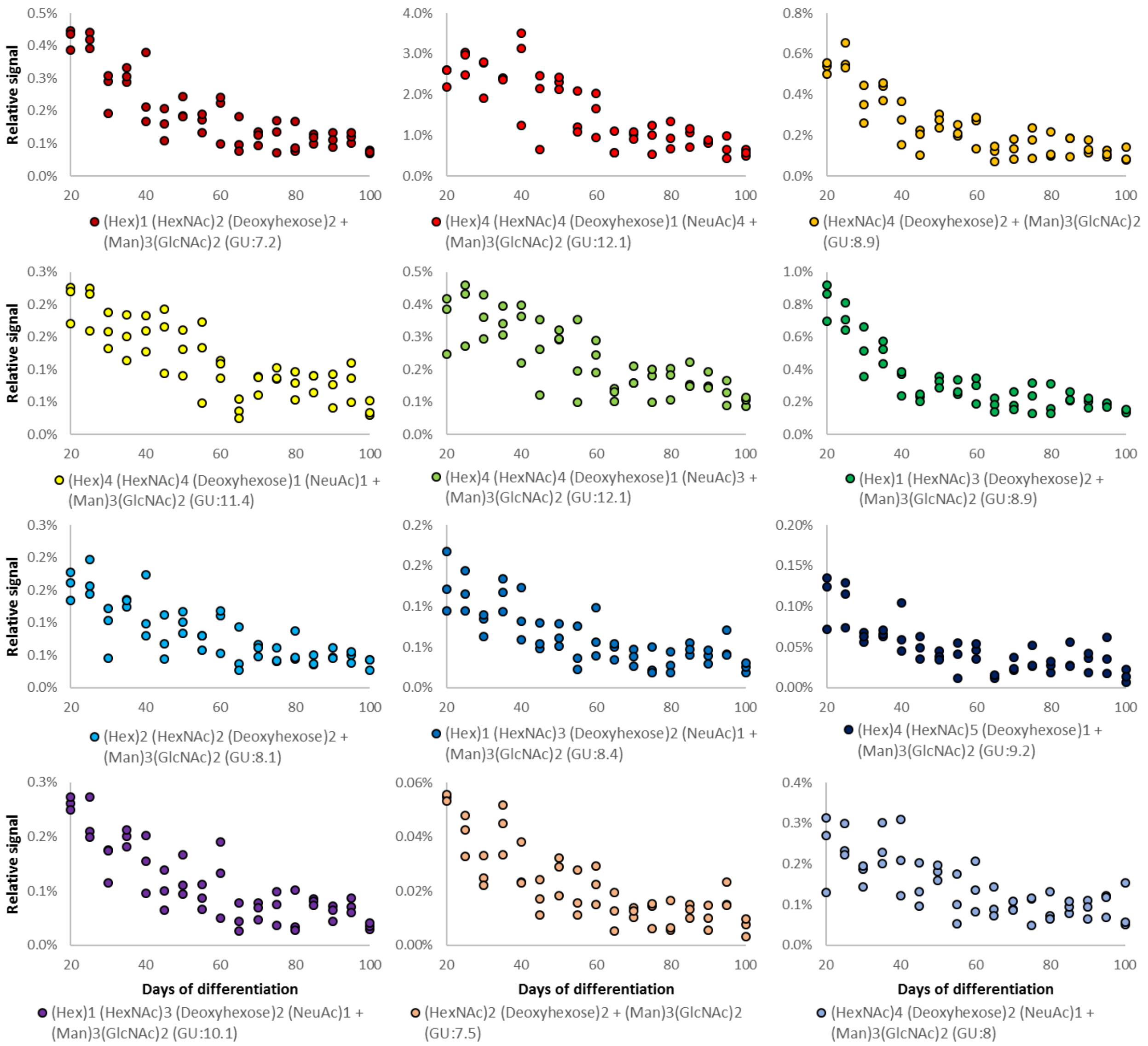


**Figure S7.** Scatter plot of relative signal for *N*-glycans from hiPSC-CM significantly negatively correlated (<-0.7) to days of differentiation

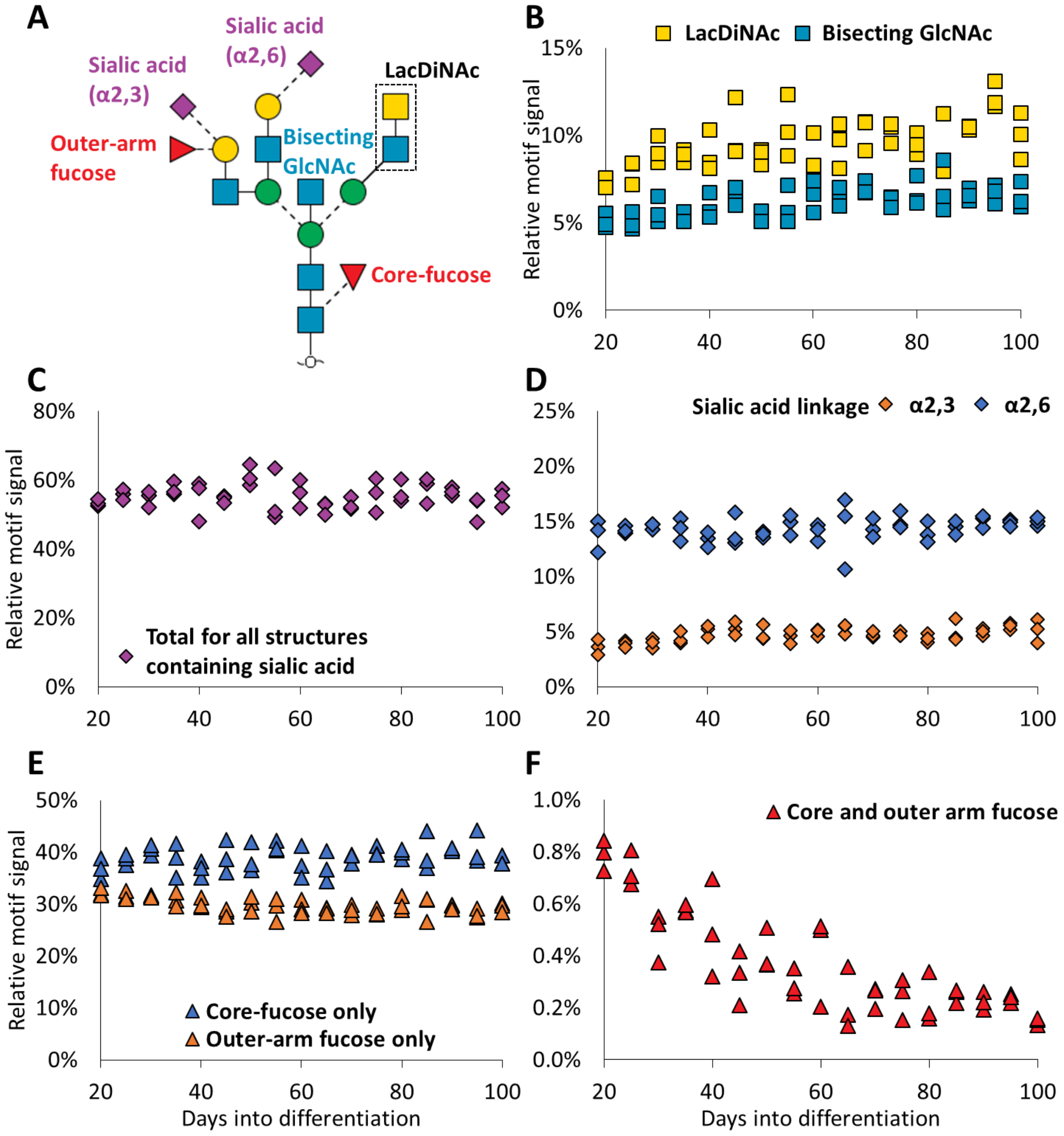


**Figure S8.** A majority of glycan structural motifs do not significantly change among hiPSC-CM collected throughout 100 days of differentiation. **A** Model *N*-glycan structure with motifs mapped to structure. **B-F** Scatter plots of relative signal for *N*-glycan structural motifs shown in A.

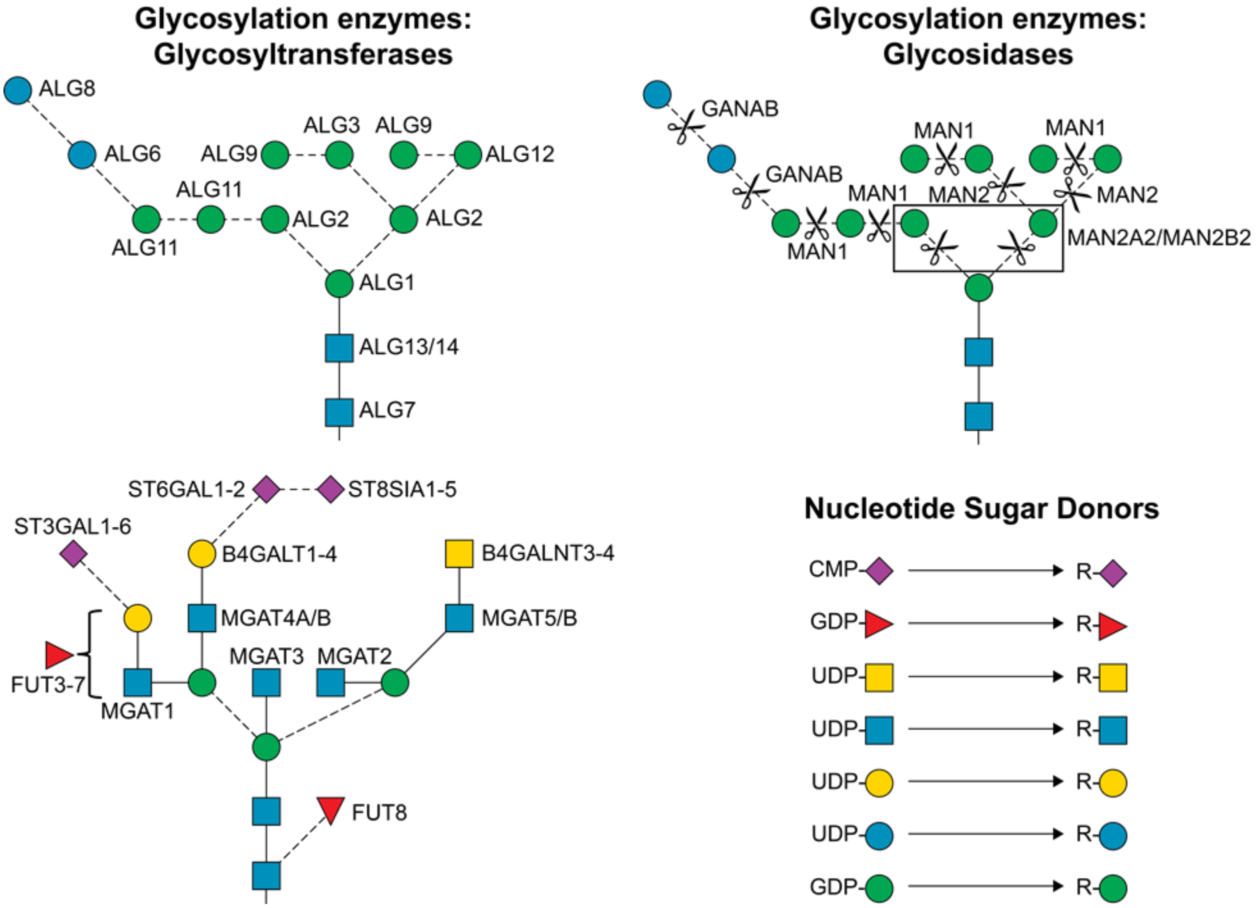


**Figure S9.** Map of the enzymes and metabolites responsible for identified and quantified glycan structures of homogenized heart tissue, primary CM and hiPSC-CM

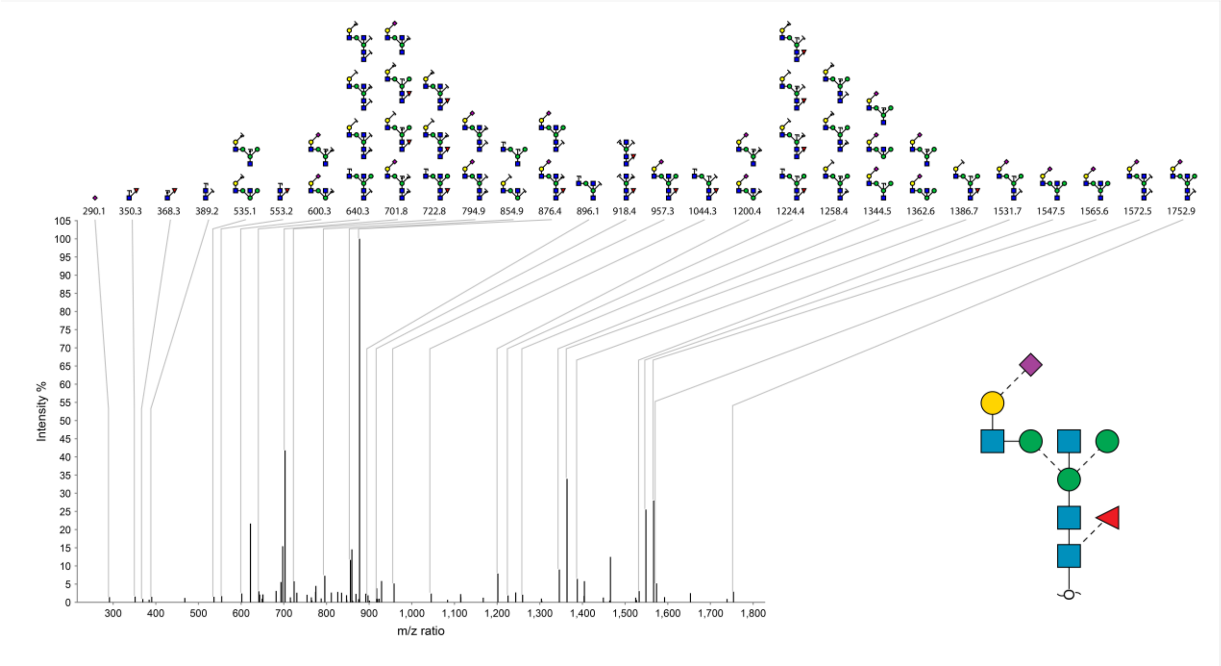
290.1 350.3 368.3 389.2 535.1 553.2 600.3 640.3 701.8 722.8 794.9 854.9 876.4 896.1 918.4 957.3 1044.3 1200.4 1224.4 1258.4 1344.5 1362.6 1386.7 1531.7 1547.5 1565.6 1572.5 1752.9
Intensity %
m/z ratio

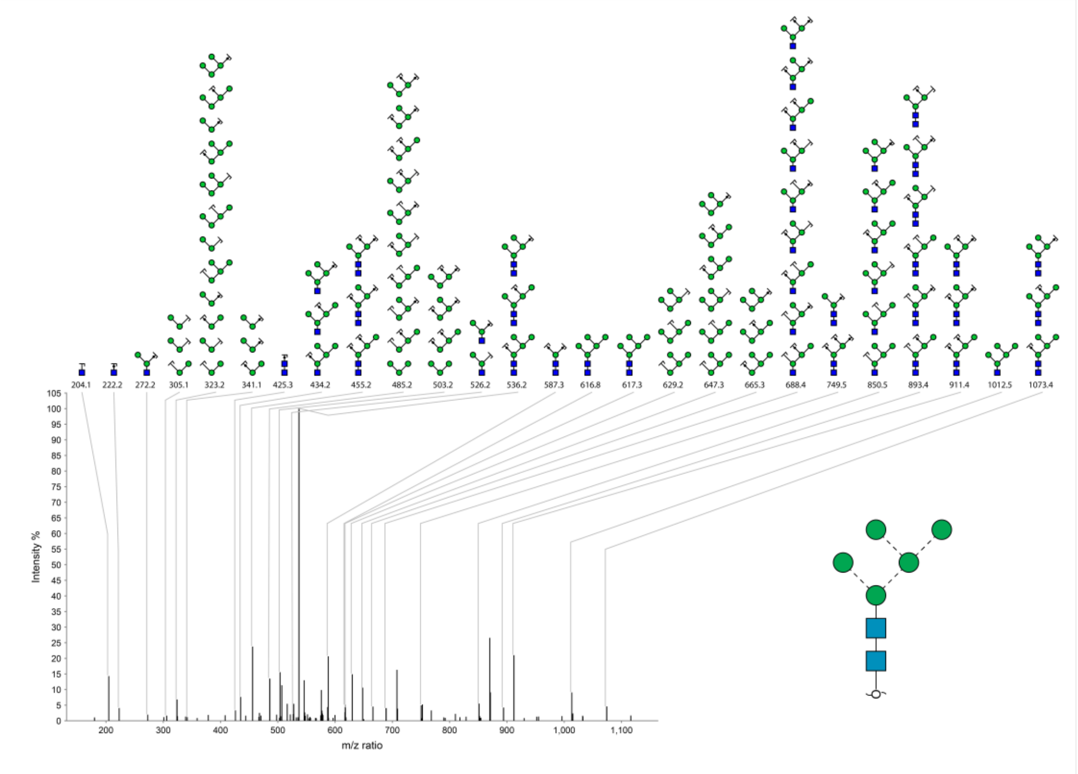
204.1 222.2 272.2 305.1 323.2 341.1 425.3 434.2 455.2 485.2 503.2 526.2 536.2 587.3 616.8 617.3 629.2 647.3 665.3 688.4 749.5 850.5 893.4 911.4 1012.5 1073.4
Intensity %
m/z ratio

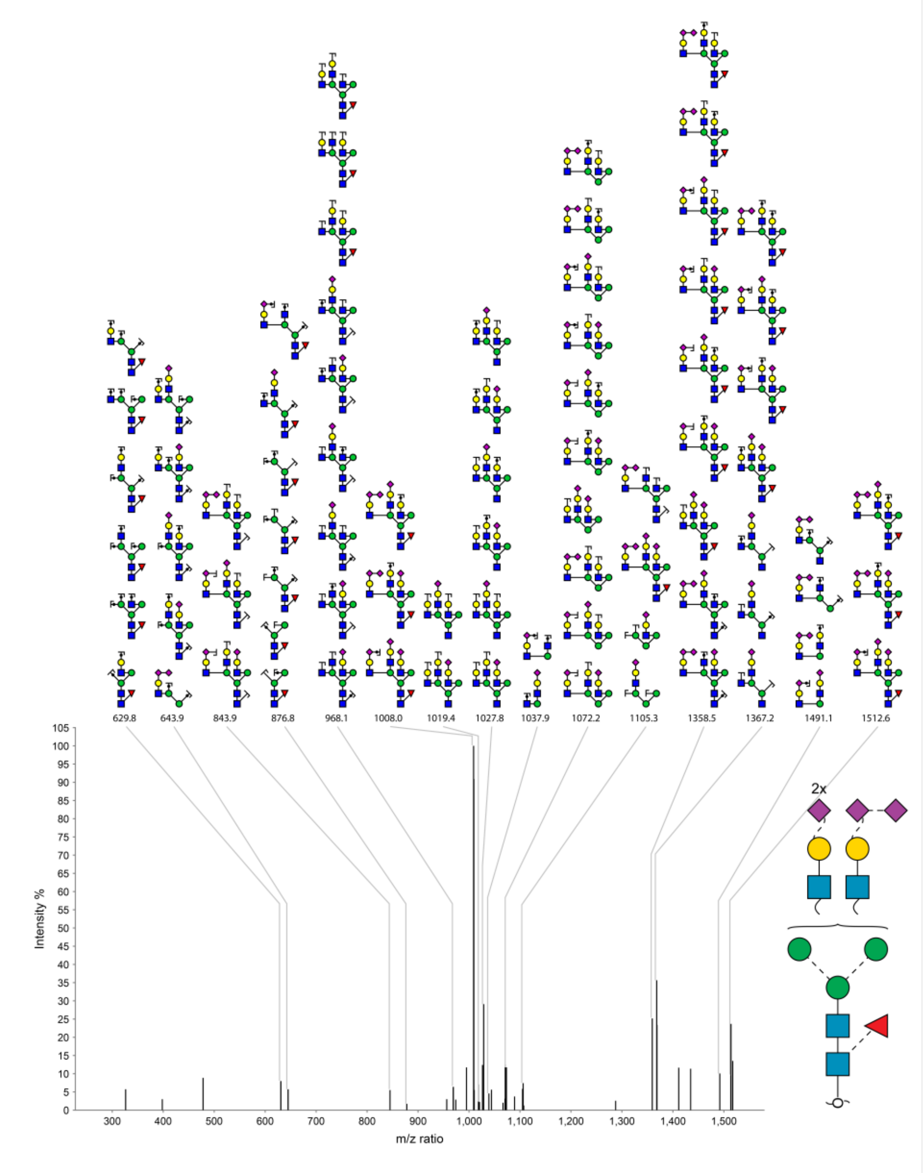


**Figure S10.** Example annotated MS2 spectra for several glycan structures in Figures 4 and 5.
